# Haplotype resolved chromosome level genome assembly of *Citrus australis* reveals disease resistance and other citrus specific genes

**DOI:** 10.1101/2022.12.20.521315

**Authors:** Upuli Nakandala, Ardashir Kharabian Masouleh, Malcolm W. Smith, Agnelo Furtado, Patrick Mason, Lena Constantin, Robert J. Henry

## Abstract

Recent advances in genome sequencing and assembly techniques have made it possible to achieve chromosome level reference genomes for citrus. Relatively few genomes have been anchored at the chromosome level and/or are haplotype phased, with the available genomes of varying accuracy and completeness. We now report a phased high-quality chromosome level genome assembly for an Australian native citrus species; *Citrus australis* (round lime) using highly accurate PacBio HiFi long reads, complemented with Hi-C scaffolding. Hifiasm with Hi-C integrated assembly resulted in a 331 Mb genome of *C. australis* with two haplotypes of nine pseudochromosomes with an N50 of 36.3 Mb and 98.8% genome assembly completeness (BUSCO). Repeat analysis showed that more than 50% of the genome contained interspersed repeats. Among them, LTR elements were the predominant type (21.0%), of which LTR Gypsy (9.8 %) and LTR copia (7.7 %) elements were the most abundant repeats. A total of 29,464 genes and 32,009 transcripts were identified in the genome. Of these, 28,222 CDS (25,753 genes) had BLAST hits and 21,401 CDS (75.8%) were annotated with at least one GO term. Citrus specific genes for antimicrobial peptides, defense, volatile compounds and acidity regulation were identified. This chromosome scale, and haplotype resolved *C. australis* genome will facilitate the study of important genes for citrus breeding and will also allow the enhanced definition of the evolutionary relationships between wild and domesticated citrus species.

## INTRODUCTION

Citrus is one of the most valuable fruit crops in the world and is widely grown in more than 100 countries under tropical, subtropical, and Mediterranean climatic conditions [1]. There are six citrus species, all of which are limes, that are native to Australia. One species is endemic to the Northern Territory whilst the other five species are primarily found in Queensland. Other populations of these species are found in New South Wales and South Australia [2]. *Citrus australis* is a slow growing, hardy plant which is naturally found in southeast Queensland [3]. *C. australis* is commonly known as Australian round lime, Dooja, Gympie Lime or Native lime. Characteristically, the trees are moderately frost tolerant, the fruits are globose or subglobose with pulp vesicles bearing large masses of oil and the seeds are monoembryonic. Although the raw fruits can be eaten, the fruits are more suitable to be used for the preparation of sauces, jams, cordials and as a flavouring agent [3].

Huanlongbing (HLB) or greening disease is caused by a vector-transmitted bacteria (*Candidatus Liberibacter*) leading to severe economic losses to the citrus industry around the world [4]. Most commercial citrus are known to be susceptible to HLB [5]. However, *C. australis* and other Australian native lime species including *Citrus australasica, Citrus glauca* and *Citrus inodora* and their derived hybrids have shown different degrees of resistance to HLB, providing highly valuable genetic resources in breeding HLB resistant cultivars, and for use as rootstocks or interstocks [6]. Recently, a novel class of small antimicrobial peptides (SAMPs) were isolated from *C. australasica* and other close relatives which can suppress the growth of HLB causing bacteria and promote host immunity in citrus [7]. However, SAMPs have not yet been identified in *C. australis* or other Australian wild limes which are resistant to HLB. The identification and characterization of genetic loci encoding the peptides and those conferring resistance against HLB is very important for the breeding of resistant citrus. Complete, high-quality genomes of resistant species will, therefore, provide enormous benefits in developing resistant cultivars.

High-quality reference genomes are a key resource for plant breeding, providing highly accurate prediction of genes in plants and supporting gene discovery [8]. PacBio circular consensus sequencing which generates Pacbio HiFi long reads (≥ 15 kb) with high base level accuracy (99.9%) outperforms the short reads and earlier long read technologies [Pacbio and Oxford Nanopore technology (ONT)] in assembling more contiguous genomes [9-11]. Hi-C based genomic scaffolding technologies have now enabled the generation of haplotype resolved chromosome level genomes in combination with Pacbio Hifi data [12]. Haplotype phasing which is still in its infancy provides unprecedented genomic resources to capture the structural variations of individual haplotypes and reveal haplotype specific variations regulating important traits. The loss of haplotype specific information in consensus genome assemblies limits their utility of informing breeding operations in highly heterozygous species [13].

Several citrus species have been sequenced and assembled over the past few years including two species (*Citrus limon* and *Citrus sinensis*) that have been anchored at the chromosome level and are haplotype resolved [14,15]. However, reference genomes of Australian limes have not yet been reported. Here we present the *de novo* chromosome-scale haplotype resolved genome assembly and annotation of genes of *C. australis* which will be a valuable resource for citrus improvement through genomic-assisted breeding approaches.

## Results

### Hifiasm assembly

The two PacBio SMRT cells yielded 30.6 Gb (90X) and 27.8 Mb (81X) of Hifi reads with Q32 median read quality (Supplementary Table S1). The contig assembly generated from HiFi reads with default parameters (default option) produced a phased assembly with two phased haplotypes. BUSCO analysis revealed that the collapsed assembly covered 98.8% universal single copy genes with an N50 of 29.5 Mb. The assembly contained 4678 contigs with a total length of 485 Mb. The two phased haplotypes; hap1 and hap2 contain a total of 4410 and 1401 contigs respectively. The total lengths of the two phased haplotypes were 470 Mb and 380 Mb. Hap1 covered 95.1% of the single copy orthologs with an N50 of 29.4 Mb, whilst hap2 covered 96.7% single copy orthologs with an N50 of 27.1 Mb (Table 1).

**Table 1.**
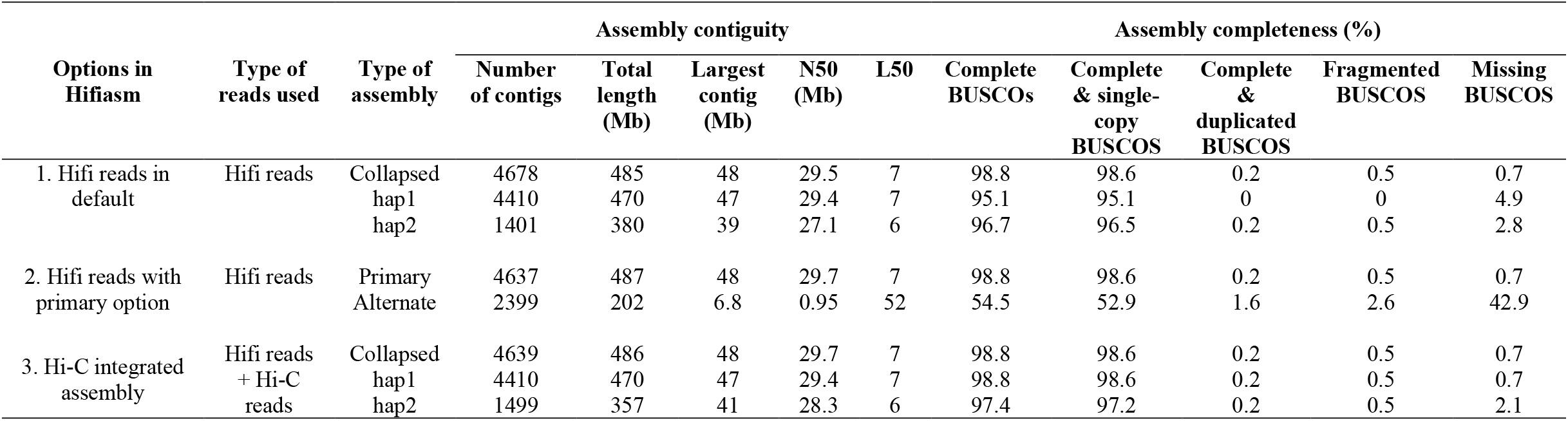
Contiguity and completeness of three *C. australis* contig level assemblies generated by Hifiasm

The primary/alternate mode generated a primary and an alternate assembly with HiFi reads. The primary assembly is comprised of 4637 contigs with 487 Mb assembly size. The BUSCO revealed 98.8% single copy orthologs for the primary assembly and the N50 was 29.7 Mb which was slightly higher than the contig assembly generated with default parameters. The alternate assembly was composed of alternate contigs that were discarded in the primary assembly. The alternate assembly was highly fragmented (N50 is 0.95 Mb) and did not cover most of the universal single copy genes (complete BUSCOs = 54.5%, missing BUSCOs = 42.9%) making it is less useful for further analysis (Table 1).

Hifiasm generated a Hi-C integrated assembly, comprising of a collapsed and a pair of phased assemblies with paired end Hi-C reads in Hi-C mode. The collapsed/consensus assembly was made up of 4639 contigs with a total length of 486 Mb. It covered 98.8% complete BUSCOs with an N50 of 29.7 Mb. The assembly contiguity and completeness in the Hi-C mode were similar to the primary assembly and the contiguity was higher than that of the collapsed assembly generated from Hifiasm default option. The two phased haplotype assemblies generated from this option; hap1 and hap2 have 4410 and 1499 contigs respectively. The two haplotype assemblies covered 98.8% and 97.4% complete BUSCOs and have 29.4 Mb and 28.3 Mb N50s respectively. Assembly statistics of two individual haplotype assemblies revealed that the contiguity and completeness of Hi-C integrated assembly are greater in comparison to the Hifi reads only assembly (Table 1).

The collapsed assembly from Hi-C mode can be characterized in four groups. There are 10 contigs greater than 13 Mb in length and 9 contigs greater than 1 Mb in length. The assembly had 36 contigs greater than 0.1 Mb and 4584 contigs less than 0.1 Mb. The dotplot analysis showed a lower sequence similarity between the phased haplotypes generated from Hifi reads only assembly (Fig. 1a), however a higher similarity between the phased haplotypes produced from the Hi-C partition options (Fig. 1b). Based on the assembly statistics, it is clear that the Hifiasm outputs the best contig assembly with the Hi-C partition options for phased haplotypes.

**Fig 1.**
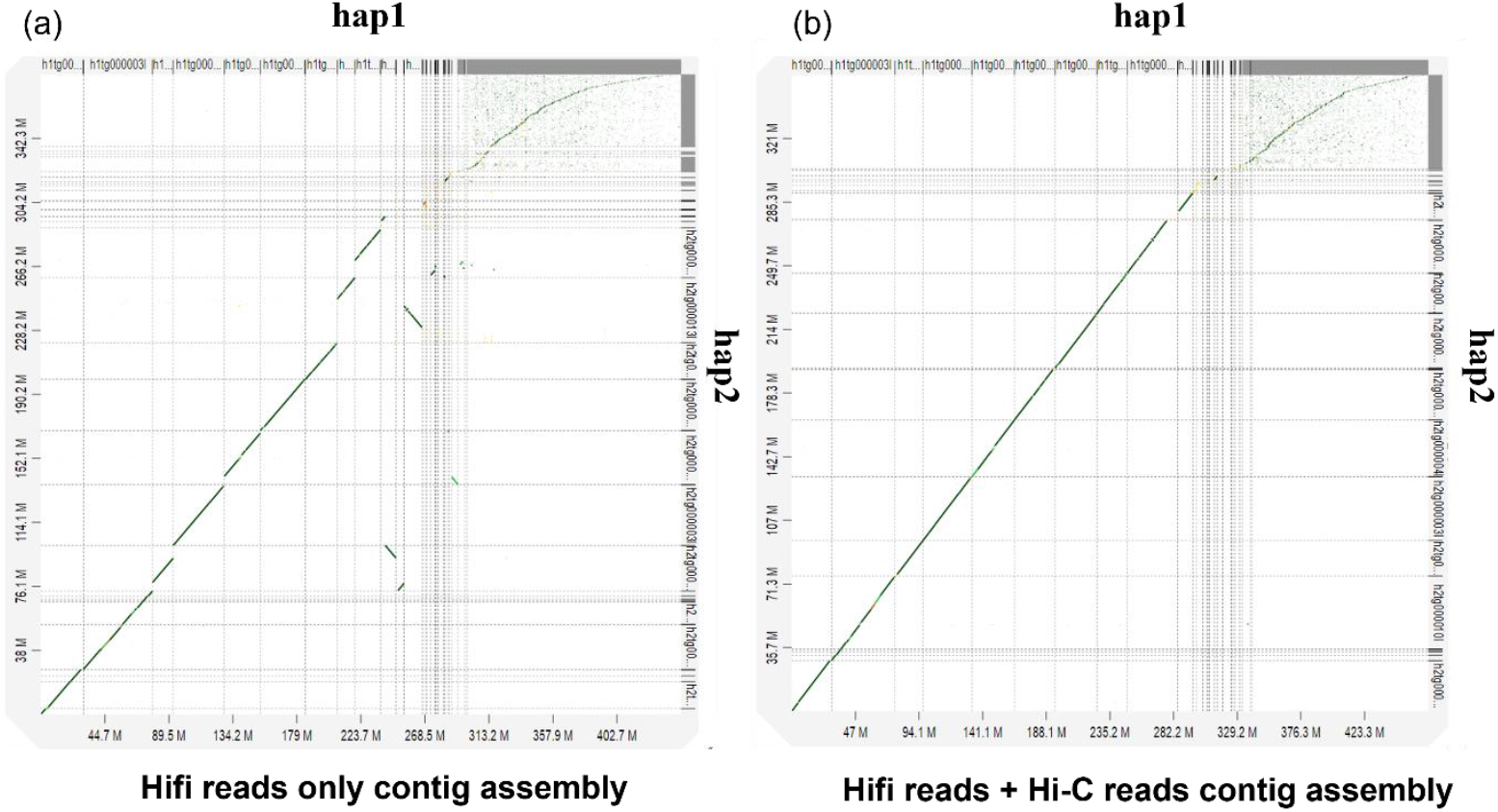
Phased haplotypes generated from two different options in Hifiasm (a) hap1 vs hap2 generated only with Hifi reads in default. The sequence similarity was less between the two phased haplotypes. (b) hap1 vs hap2 generated with Hifi reads and Hi-C reads. The two phased haplotypes had high sequence similarities

### Chromosomal scale pseudochromosome generation using Hi-C data

The three contig assemblies was subjected to scaffolding to further understand assembly contiguity and completeness at the scaffold level. Hi-C proximity ligation libraries produced in two lanes generated a total of 656 M paired-end reads. Hi-C scaffolding of the first option (Hifi reads in default) generated 4663 scaffolds with 29.7 Mb N50 and 98.8% of complete BUSCOs. In the second option, the primary assembly generated 4618 scaffolds with 31.3 Mb N50 and 98.8% complete BUSCOs. The third option (Hi-C mode) generated 4642 total number of scaffolds with 31.3 Mb and 98.8% complete BUSCOs (Supplementary Table S2). Hi-C scaffolding revealed that the second and third options generated assemblies with similar contiguity which outperformed the first option where only Hifi reads were used in default.

Based on the assembly statistics and the ability to generate phased assemblies, we selected the Hi-C integrated Hifiasm assembly as the best assembly to be used for subsequent downstream analysis. The dotplot between Hi-C scaffolds against Hi-C integrated HiFiasm contigs assembly is shown in Fig. 2a. The 4642 scaffolds could be characterized into three subgroups including large scaffolds (from 24 Mb - 48 Mb), medium sized scaffolds (from 1.5 Mb - 4.5 Mb) and small sized scaffolds (less than 1.5 Mb). The scaffolding generated nine main pseudomolecules corresponding to the nine chromosomes with a total length of 311Mb, an N50 of 36.3 Mb and 98.8% completeness of conserved single copy orthologs. The pseudochromosome numbers were assigned to the corresponding scaffolds based on synteny between the present assembly of *C. australis* and other three genomes: *C. sinensis* (sweet orange) (Fig. 3a), *C. maxima* (pummelo) (Fig. 3b) and *C. limon* (lemon) (Fig. 3c).

**Fig 2.**
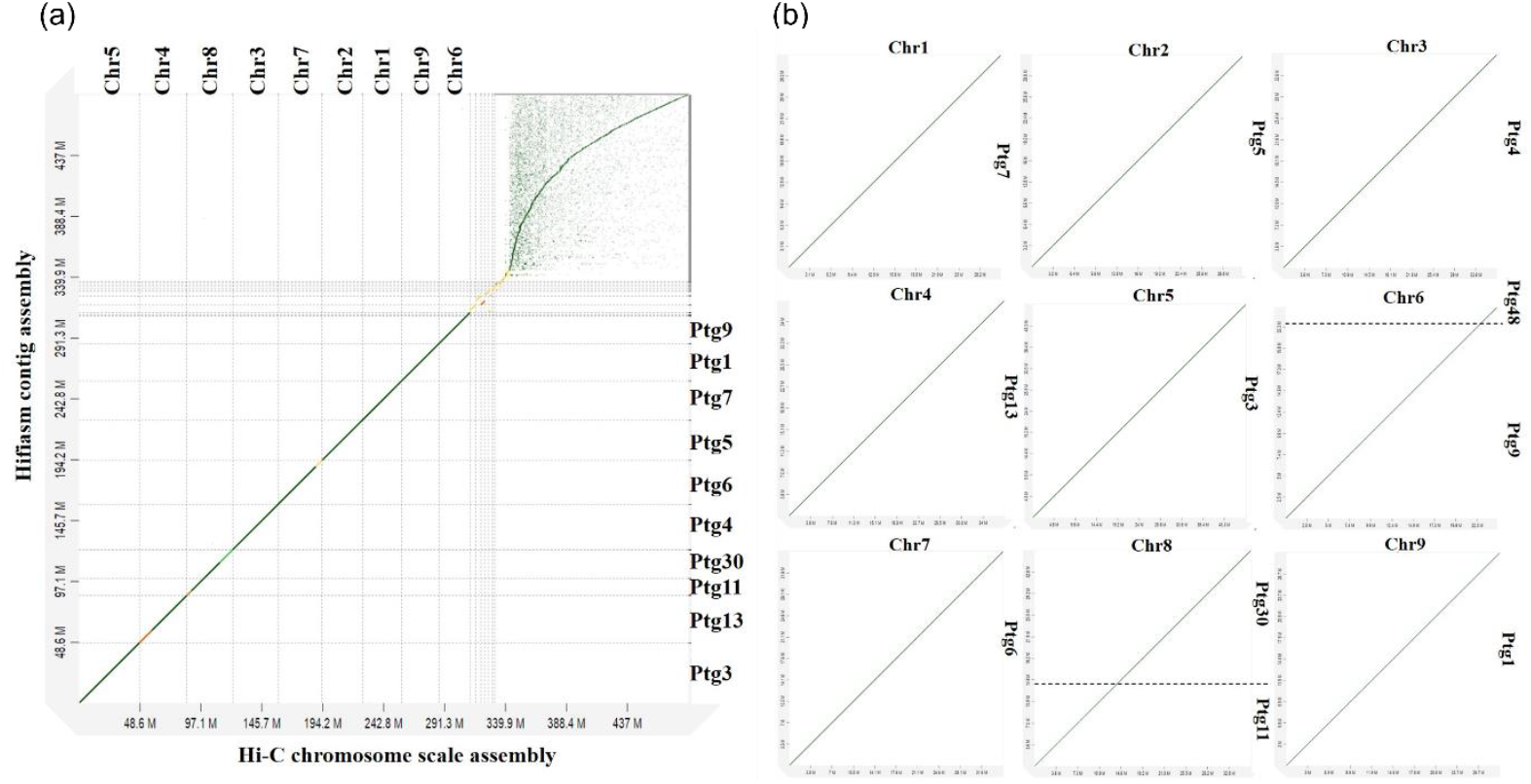
Hi-C chromosome scale pseudomolecules and the corresponding Hifiasm contigs. (a) The x-axis shows the chromosome scale assembly generated with Hi-C data. The y-axis indicates the contig assembly generated by Hifiasm with Hifi data and Hi-C paired end data. The lengthiest nine scaffolds are named with corresponding chromosome numbers based on the synteny analysis with previously published genomes. The middle set of scaffolds with the lengths of 1.5 Mb – 4.5 Mb contain large clusters of unknown repeats and are not assembled into 9 chromosomes. Small scaffolds with lengths less than 0.1 Mb are chloroplast genome fragments inserted within the nuclear genome and are shown at the top right corner of the image. (b) Individual chromosomes and corresponding Hifiasm contigs. Chromosomes 6 and 8 are covered by two contigs whereas the other chromosomes are covered by one contig.

**Fig 3.**
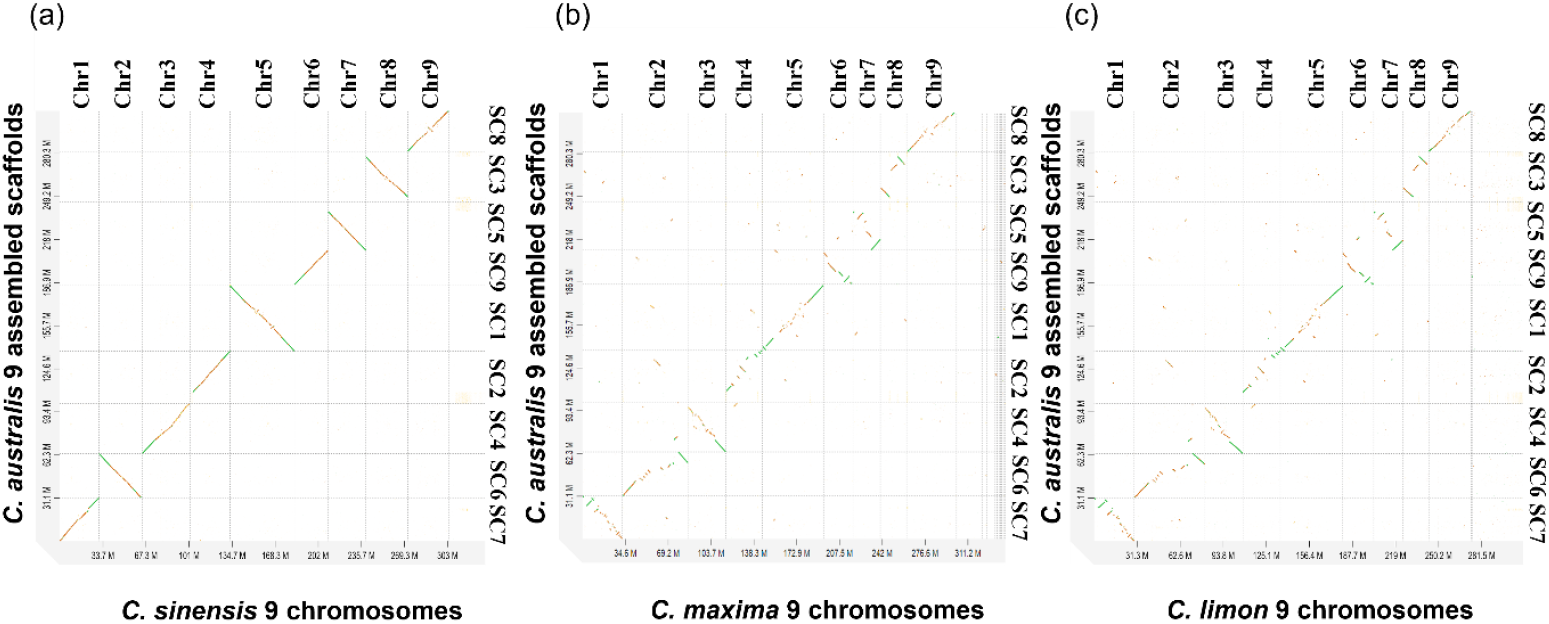
Dotplot analysis showing the synteny between *C. australis* assembled nine pseudomolecules and nine chromosomes of other three citrus genomes a, *C. sinensis;* b, *C. maxima;* and c, *C. limon*. The sequence similarities between the corresponding pseudomolecules were used to rename *C. australis* scaffolds. Chr1, Chr2, Chr3, Chr4, Chr5, Chr6, Chr7, Chr8, and Chr9 correspond to scaffolds 7,6,4,2,1,9,5,3,8

Out of the nine pseudochromosomes, seven (Chr1, Chr2, Chr3, Chr4, Chr 5, Chr7, Chr9) were covered by one single HiFi contig and two (Chr6 and Chr8) were represented by two Hifi contigs (Fig. 2b, Supplementary Table S3). The nine pseudochromosomes were in the range between 24.8 Mb – 48.1 Mb, where chromosome five was the longest (48.1 Mb) and chromosome six was the shortest (24.8 Mb). Among the nine pseudomolecules, pseudomolecules one, three, five and seven had telomere repeats at both ends representing full chromosomes while the other pseudomolecules had telomere repeats only at one end. Pseudomolecules six and eight were spanned by two contigs and only one peripheral chromosomal region of the two pseudomolecules had telomeres (Supplementary Table S3). There were also five medium sized scaffolds which did not belong to the nine chromosomes ranging from 1.5 Mb – 4.5 Mb (Figure 2a). The scaffolds of individual haplotypes were assigned with chromosome numbers corresponding with the nine pseudo chromosomes of *C. australis* collapsed assembly based on dotplots (Fig. 4). Pseudo chromosome five was the largest of the two haplotypes (hap1 - 46.9 Mb, hap2 – 47.4 Mb) and the pseudochromosome 6 was the smallest (hap1 – 24.4 Mb, 24.6 Mb).

**Fig 4.**
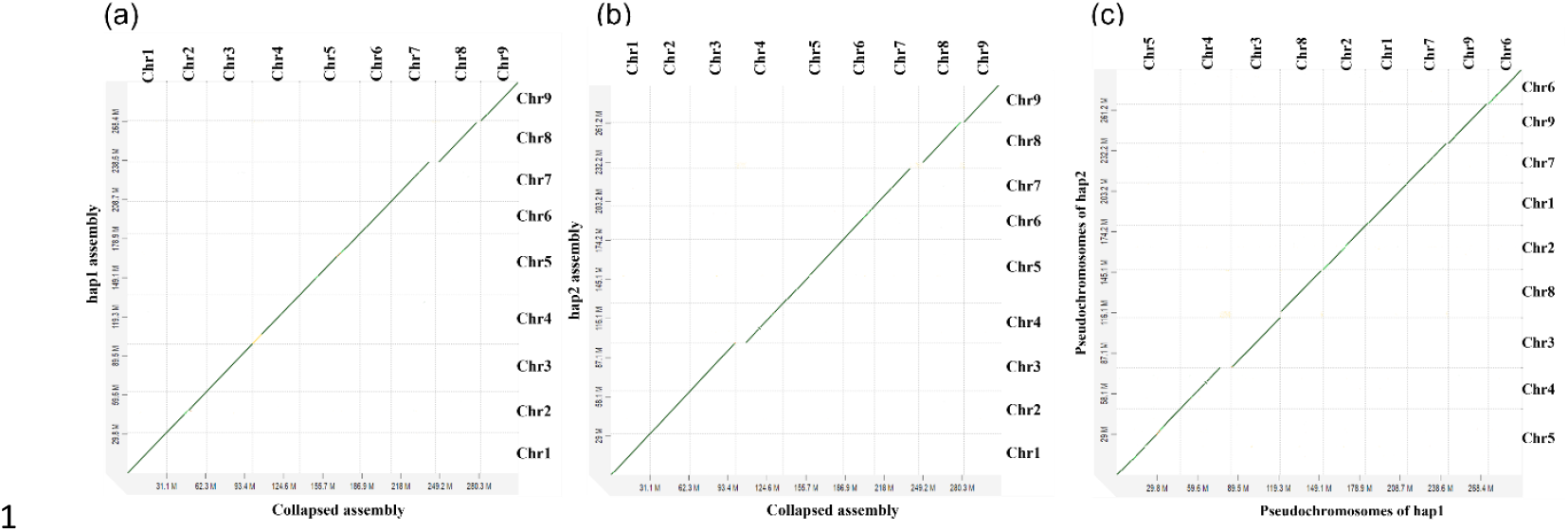
Pseudochromosome scale collapsed assembly vs pseudochromosome scale haplotype 1 and haplotype 2 assemblies of *C. australis*. (a) Collapsed assembly vs hap1 assembly (b) Collapsed assembly vs hap2 assembly (c) hap1 assembly vs hap2 assembly

### Organelle genome analysis

Alignments of the complete *C. australis* chloroplast genome (Supplementary Fig. S1) with the small sized scaffolds showed that large parts of the scaffolds 27-4642 showed high similarities with the chloroplast genome (Supplementary Fig. S2c). Among the medium sized scaffolds, only scaffolds 12 and 13 contain small fragments of the chloroplast genome (Supplementary Fig. S2b). Sequence similarities with some parts of the top 9 scaffolds with the chloroplast genome indicate the insertion of chloroplast sequences within the nuclear genome (Supplementary Fig. S2a).

### Comparisons of assembly size with k-mer approaches and flow cytometry estimates

Genome estimates varied depending on the k-mer value employed in the analysis. K-mer 21 was used for genomescope in one approach as it was the most widely executed k-value in other research [14,16] and the recommended length by Genemoscope based on the computational accuracy and speed [17]. The k-mer depth distribution histogram was a bimodal profile which is the typical nature of heterozygous genomes with a short peak around 75X coverage and high peak around 150X coverage (Supplementary Fig. S3a). The estimated genome size with k-mer 21 was 297 bp with 0.503% heterozygosity. The kmergenie approach generated abundance histograms for different values of k ranging from k=17-121 and predicts the best k value as 97. The predicted best k=97 was then used for genome size estimation using genomescope which dramatically increased the genome size to 318 bp (Supplementary Fig. S3b). The results indicated that higher values of k resulted in higher genome sizes in genomescope. The genome size was estimated to be 340.5 Mb ± 0.5416 % CV using flow cytometry (Supplementary Fig. S4). The estimated genome assembly size of the nuclear DNA content was 330.6 Mb.

### Repeat identification and masking

Nine pseudochromosomes of the collapsed assembly were annotated for repeats and genes as nine chromosomes had the same number of BUSCOs as the whole genome (98.8%). A large portion of the genome (52%) was comprised of interspersed repeats where the majority of them were unclassified. Among the classified transposable elements, LTR elements were the predominant type (21.0%). Among them, 21,192 regions were covered by LTR Gypsy elements accounting for the highest total size of the repetitive regions (30,530,861 bp) (9.8 %) in the genome, followed by LTR Copia (23,997,689 bp) (7.7 %). In addition to these dominant LTR elements, other LTR elements such as caulimovirus, ERV1, Ngaro, and pao were scattered throughout the genome with varied sizes. The total number of regions associated with Long Interspersed Nuclear Elements (LINEs) was 6205 accounting for a total of 5,269,998 bp (1.66 %). There were no SINEs in the present genome. DNA transposons were present in smaller proportions (3.4%). The major types of DNA transposons were DNA/MULE-MuDR (1.3%) and DNA/hAT-Ac (0.8%). In addition to the transposable elements, a small proportion was composed of simple repeats (0.96), low complexity repeats (0.22%), small RNA repeats (0.1%) and satellites (0.01%). The highest number of repetitive regions were annotated in Chr5 24,034,718 (7.7%), which is the longest chromosome, and the least was recorded in Chr6, which is the shortest chromosome (3.92) (Table 02, Fig. 5, Supplementary Table S4; S5).

**Table 02.**
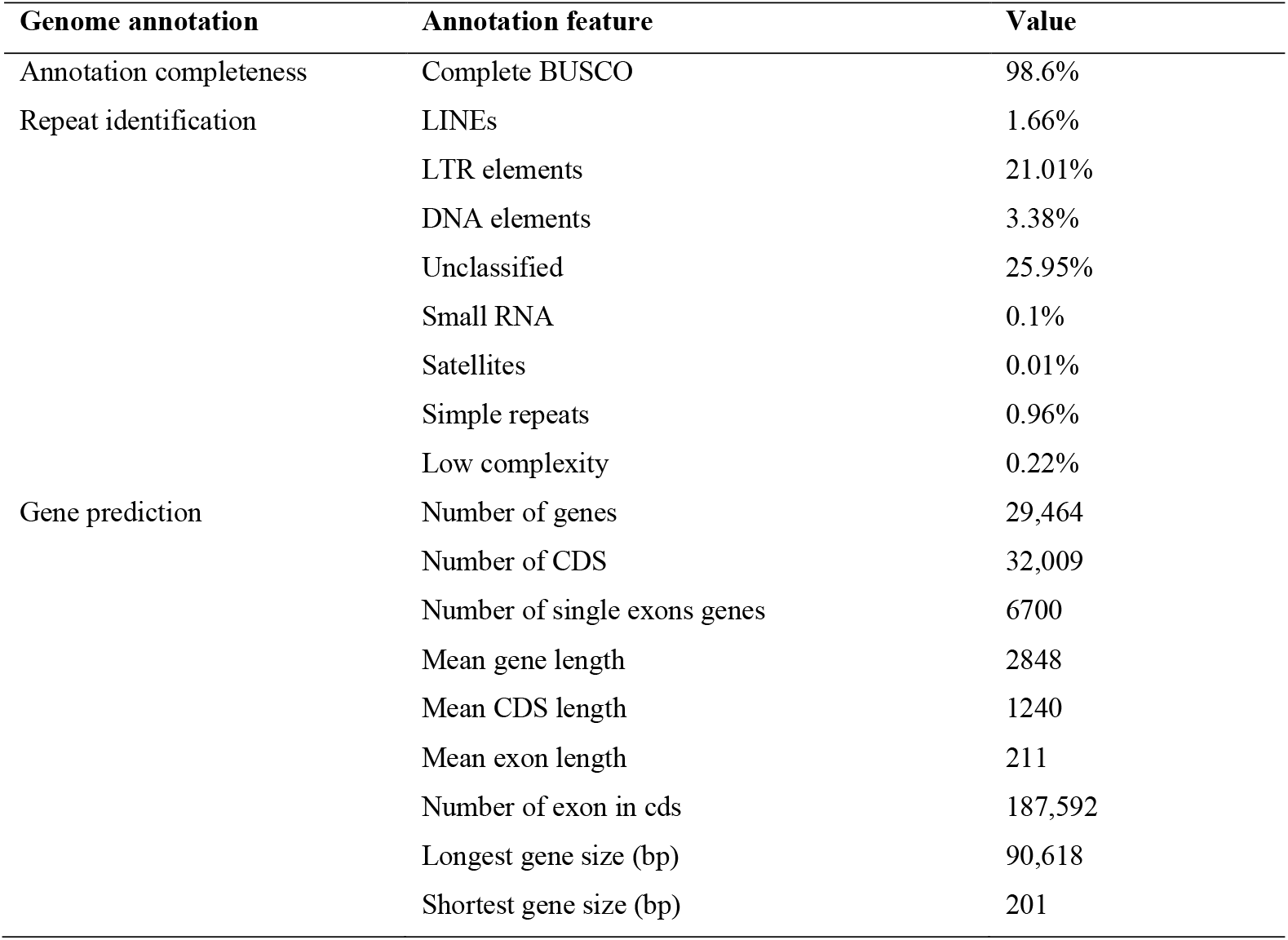
Summary of BUSCO, repeat identification and gene prediction statistics for the collapsed genome

**Fig 5.**
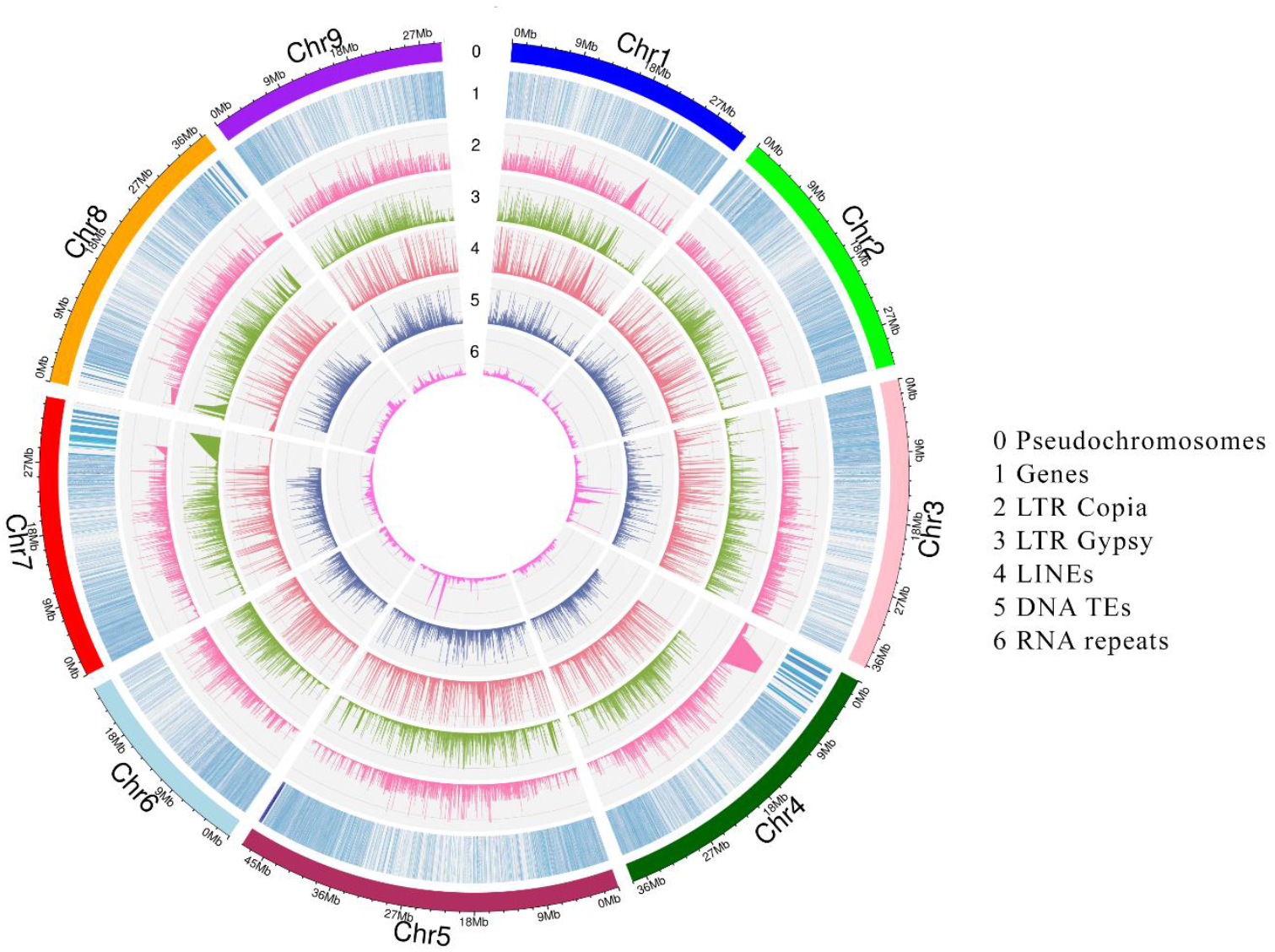
Characterization of repetitive regions and genes in *C. australis* genome. (0) Nine pseudochromosomes (Mb), (1) Regions of predicted genes (2) Regions of LTR Copia elements, (3) Regions of LTR Gypsy elements, (4) Regions of LINEs, (5) Regions of DNA TEs, (6) Regions of rRNA, tRNA and snRNA repeat regions

A total of 165 Mbp (53.17%) of the genome was masked by repeat masking software. Hard masking and soft masking masked 165, 632, 936 bp (53.17%) of the genome. The hard masking with -nolow flag causes the software to hard mask all the repetitive regions excluding low complexity DNA such as Poly-purine or poly-pyrimidine stretches, or regions of extremely high AT or GC content and simple repeats accounting for 162, 287, 041 bp (52.10%) of the genome [18].

### RNA-seq read alignment

A total of 37.6 Gb (X110) in 250 million paired-end RNA-seq reads were mapped to the genome. Quality trimmed only RNA-seq reads and Quality and adapter trimmed RNA-seq reads were used for mapping to understand the effect of adapter trimming on overall alignment rates. The overall alignment rates for the quality trimmed only RNA-seq data with the unmasked, softmasked, hard masked with nolow option, and hard masked genomes were 60.6%, 60.6%, 52.2%, 50.8% respectively. The overall alignment rates of the quality and adapter trimmed RNA-seq data were improved in all the cases that were 79.1%, 79.1%, 68.1%, and 66.5% respectively.

### Gene prediction

Gene prediction was done for the unmasked and masked genomes to understand the impact of repeat masking on gene prediction. Quality trimmed only and quality and adapter trimmed RNA-seq data were independently used for gene prediction. Higher number of genes were predicted with quality and adapter trimmed RNA-seq data whereas low number of genes were predicted with quality trimmed only RNA-seq evidence (Supplementary Table S6). The higher number of genes is due to the higher alignment rates of quality and adapter trimmed RNA-seq data with the genome. The predicted number of genes with quality and adapter trimmed RNA-seq evidence for the soft masked genome was 29,464 (Figure 5, Table 02). In the genome, some protein coding genes occupy repeat regions which will not be counted during the gene prediction with hard masking. Braker can still assess the gene sequences in repeat regions in a soft masked genome. Due to this, the softmasked genome is preferred over the hard masked genome for gene prediction. The highest number of genes was recorded in chromosome 5 (5240) while the lowest was recorded in chromosome 6 (2346) (Table 3). 98.6% of complete BUSCOs indicate the high completeness of the protein coding gene prediction (Table 02).

**Table 3.**
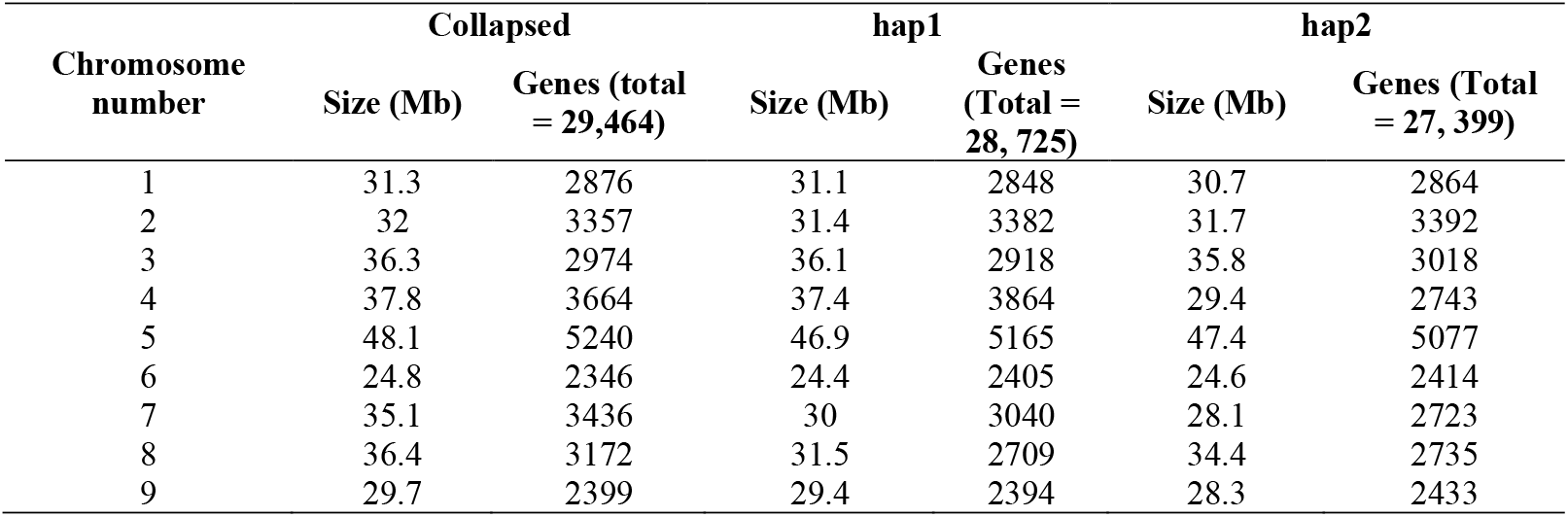
Sizes of the pseudochromosomes of collapsed genome and two haplotypes and the numbers of genes in each genome

The total number of genes predicted for hap1 was 28725 and for hap2 was 27399. The total lengths of the haplotype assemblies (nine pseudochromosomes) differed by 7.8 Mb and the total number of genes differed by 1326. The size of the hap1 is 13.3 Mb shorter than the collapsed assembly and the total number of genes differed by 739 between the collapsed and hap1 genomes. The size of the hap2 was 21.1 Mb shorter than the collapsed assembly and the gene number differed by 2065 between the hap2 and collapsed genomes. (Table 3).

### Non-coding genes prediction

Barrnap tool identified a large 5S ribosomal block in chromosome 6. A small number of 5S rRNA genes were predicted in Chr3, Chr4 and Chr7. No rRNA genes were found in chromosome 2. Chr1 only had 18S rRNA genes. 5S, 5.8S, 18S rRNA genes in Chr5, 5S, 5.8S, 18S and 28S rRNA genes in Chr6 and Chr8 and 5.8S, 18S and 28S rRNA genes in Chr9 were predicted by the tool.

### Functional annotation

Of the total number of predicted CDS by Genemark trained Augustus (32,009), BLAST hits were obtained for 28,222 CDS (25,753 genes). The highest number of BLAST top hits (21, 642) were from *Citrus sinensis*. In addition, more than 8422 Top BLAST hits were from *Citrus clementina* and more than 5501 hits were from *Citrus unshiu*. A very small percentage of hits (537) were from other species (Supplementary Fig. S5A). The best hit with the smallest E value (below E 10^−9^) of all the annotations were used to describe the predicted genes. Among the transcripts with BLAST hits, 21,401 CDS (75.83%) were annotated with at least one GO term (Supplementary Fig. S5B). GO and Enzyme code distributions are given in Supplementary Fig. S5C; S7D. The coding potential assessment for BLAST hits with no BLAST hits [3808 CDS (3728 genes)] using *Arabidopsis thaliana* models revealed eight non-coding transcripts (8 genes) and 3800 coding transcripts. Based on citrus models, we identified 52 non-coding transcripts (48 genes) and 3756 coding transcripts. The predicted number of total non-coding transcripts was 54 corresponding to 50 genes from both models after removing the redundant transcripts.

### Important genes in citrus

#### Antimicrobial peptides

A novel class of relatively short stable antimicrobial peptide (SAMP) with 67 amino acids (aa) has recently been detected from two wild Australian citrus species and a few citrus relatives [7]. The BLAST search identified two homologous genes in *C. australis* encoding a stress-response A/B barrel domain-containing protein HS1 (Supplementary Fig. S6). The first gene g9664 residing in chromosome 9, has two transcripts, one encoding 153 aa (g9664.t1) with 37% sequence similarity and the other transcript encoding 192 aa (g9664.t2) with 31% similarity with the 67 SAMP sequence previously characterized from resistant species. The sequence alignment showed that a substantial portion of 67 SAMP is present as a part of these two long peptides produced by *C. australis* with few amino acid substitutions and insertions (Supplementary Fig. S7a).

Another gene, g2059 on chromosome 8, encodes a 114 aa peptide (stress-response A/B barrel domain-containing protein HS1) with 21% sequence similarity with 67 SAMP (Supplementary Fig. S7b). HLB susceptible species such as *C. clementina* and *C. sinensis* have A/B barrel domain-containing protein HS1 with long amino acid sequences with different lengths (*C. clementina* – 118 aa, and 114 aa, *C. sinensis* – 109 aa, 114 aa, 126 aa, 175 aa). The *C. clementina* and *C. sinensis* proteins (114 aa) are identical to that of *C. australis* (114 aa) encoded by g2059 gene except for one SNP indicating that this is a common antimicrobial protein (AMP) present in these plants.

#### Other defense related genes

Other defense-related genes that were annotated in *C. australis* genome, might potentially be involved in HLB resistance (Fig. 6). Genes encoding 17 guanine nucleotide-binding proteins (Supplementary Table S7), 13 pathogenesis-related proteins (Supplementary Table S8), and 76 Leucine rich repeat (LRR) proteins (Supplementary Table S9) were identified in the genome.

**Fig 6.**
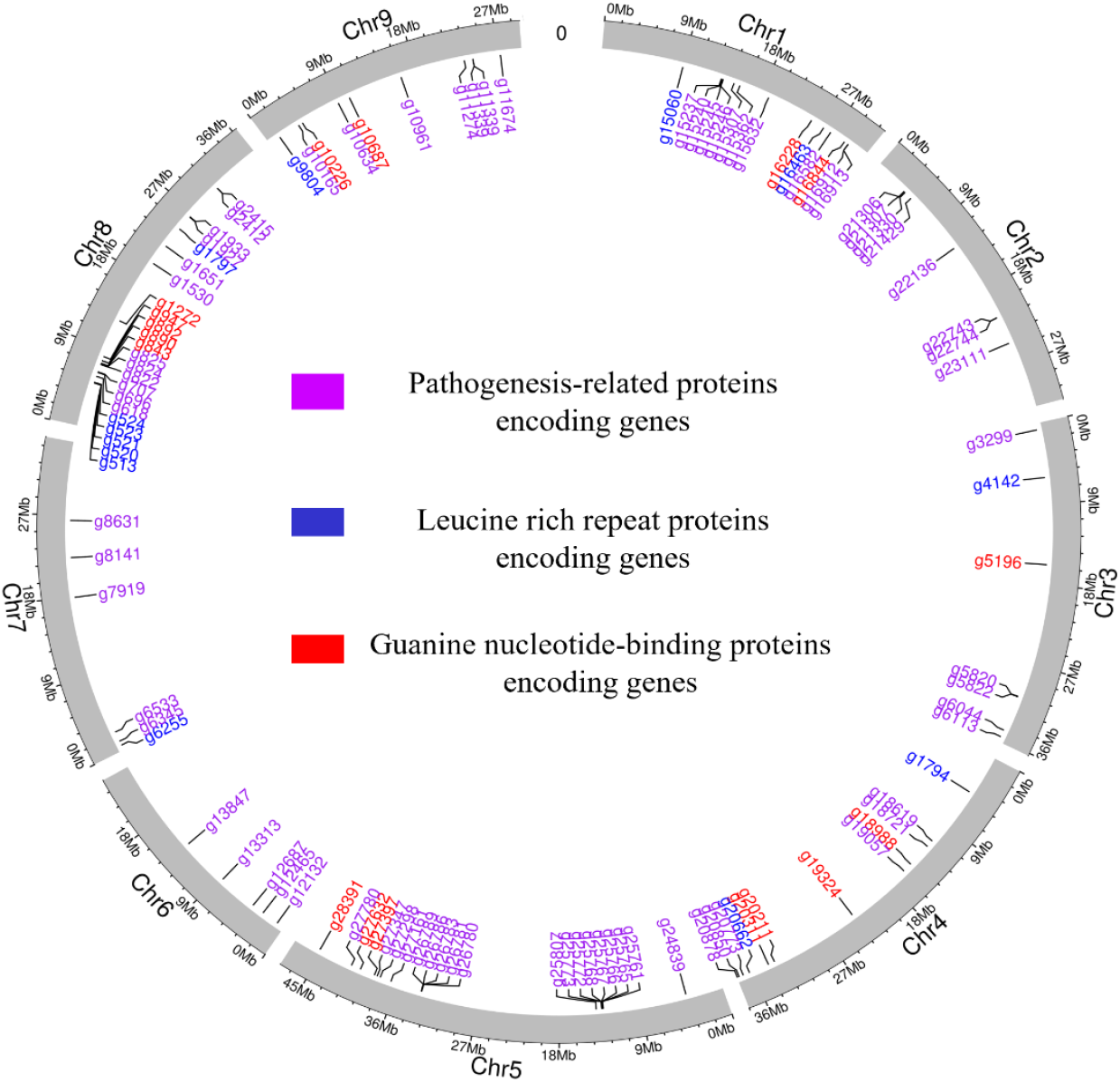
Circos plot showing the location of defense-related genes in the *C. australis* genome Seventeen guanine nucleotide-binding proteins encoding genes (red), 13 pathogenesis-related proteins encoding genes (purple), and 76 Leucine rich repeat (LRR) proteins encoding genes (blue) were identified in the genome.

#### Citrus acidity genes

We identified the homologs of two key acidity controlling genes; *PH1*(P-type Mg(2+) transporter) and *PH5* (P-type H(+)-exporting transporter) in *C. australis* genome by searching the sequence similarity with the corresponding protein sequences of *C. sinensis* for *PH1* (Cs1g20080) and *PH5* (Cs1g16150) [19] (Supplementary Table S10). The genes homologous to *PH1* and *PH5* were present on chromosomes 1 and 5 respectively. In addition to *PH5*, there were nine other genes encoding P-type H(+)-exporting transporters on chromosomes 4 and 6. PH1 protein of *C. australis* is distinguishable from other cultivated citrus species with a sweet flavor such as *C. sinensis, C. clementina* and *C. reticulata* by a few amino acid substitutions (Supplementary Fig S8). The different expression levels of *PH1* and *PH5* are thought to be regulated by the transcription factors CitAN1 (basic helix-loop-helix transcription factor family protein), CitPH3 (WRKY transcription factor 44 isoform X1) and CitPH4 (R2R3-MYB family transcription factor). *C. australis* homologs of these genes were identified using the respective protein sequences of *C. sinensis* and they were present in one copy in the genome and encode only one transcript except for PH3 which had two transcripts.

The citric acid level of juice sacs is also determined by the degradation of citric acids by a combination of enzymes including cytosol and mitochondrial aconitase (CitAco), NADP-isocitrate_dehydrogenase (CitIDH), and Glutamine synthetase (CitGS) and Glutamate decarboxylase (CitGAD). We identified three genes of aconitate hydratase protein CitAco3 (g19909 – Chr4, g22425 - Chr2, g17125 – Chr1) each encoding one transcript. Three CitIDH genes (g5386 – Chr3, g11983 – Chr9, g21560 – Chr2), three CitGS genes (g10022 – Chr9, g12494 – Chr6, g7198 – Chr7) and two GAD genes (g22832 – Chr2, g26170 – Chr5) in *C. australis* genome (Supplementary Table S10, Supplementary Fig. S6).

#### Volatile compounds synthetic genes

We identified a few key genes governing the synthesis of terpenoids in *C. australis* via the two main terpenoid biosynthetic pathways [3-hydroxy-3-methylglutaryl-CoAsynthase-2 (HMGS-2), mevalonate kinase (MVK), phosphomevalonate kinase (PMK), farnesyl pyrophosphate synthase (FPPS) in the MVA pathway and geranyl pyrophosphate synthase (GPPS) and geranylgeranyl diphosphate synthase (GGPPS) in the MEP pathway] (Supplementary Fig. S6). There was one locus on chromosome 3 encoding PMK, three loci for MVK on Chromosomes 2,3 and 5, two loci for HMGS residing on chromosomes 5 and 9, one locus for FPPS in chromosome 4, one locus for GPPS in chromosome 2 and 18 loci encoding GGPPS related genes (Supplementary Table S11). In addition, we identified 79 Terpene Synthase (TPS) genes in *C. australis* genome (Supplementary Fig. S9). Of these, 37 genes were responsible for the synthesis of monoterpenes (Beta-myrcene/(E)-beta-ocimene synthase 2, S-(+)-linalool synthase, d-limonene synthase, (E,E)-geranyllinalool synthase, tricyclene synthase, alpha-terpineol synthase, gamma-terpinene synthase), 24 genes for sesquiterpenes (alpha-humulene, alpha-copaene synthase-like) and nine genes were involved in the synthesis of diterpenoids (Ent-copalyl diphosphate synthase, cis-abienol synthase, Ent-kaur-16-ene synthase). Among the monoterpenes, eight genes encoding d-limonene synthase and ten genes encoding Beta-myrcene/(E)-beta-ocimene synthase 2 were annotated on Chr 2, 3 and 8. The highest number of genes (22) were annotated for alpha-humulene synthase which is responsible for the production of sesquiterpenes.

## Discussion

Here we present the first report of a high quality, complete, and haplotype resolved chromosome level genome assembly for *C. australis*. The size of the assembled nuclear genome was 330.6 Mb of which 311 Mb represents the nine pseudochromosomes and the remaining 19.6 Mb could not be anchored to the 9 chromosomes due to the presence of large clusters of uncharacterized repeat elements. With that the present assembly has anchored 94.1% of the total nuclear genome to the chromosome level. The N50 is higher for this genome than that for all other published citrus genomes [14,15] suggesting that it could potentially be a high-quality reference genome for limes and their hybrids. We achieved the highest assembly BUSCO completeness (98.8%) thus far in citrus and 98.6% annotation completeness which is the third largest among published genomes. Assembly statistics showed that when Hi-C reads were integrated with Hifi reads, it generated phased assemblies with slightly improved contig and scaffold N50s for the collapsed genomes and the two haplotypes. Our results reveal that the combined use of Hifi reads and Hi-C reads leveraged the maximum potential of Hifi reads in assembling heterozygous, highly repetitive plant genomes which is in agreement with previous studies [20].

Hifiasm can run in 4 modes depending on the availability of sequence data. Hifiasm (trio) mode can generate fully haplotype resolved assemblies if maternal and paternal short reads are available. Hifiasm (primary/alternate) mode generates two assemblies; one is the primary assembly containing long contigs which are not haplotigs and an alternate assembly containing haplotigs which are fragmented. Hifiasm (dual) generates two hifiasm assemblies only with Hifi reads which are not fully haplotype resolved and are more likely a primary assembly. Hifiasm (Hi-C) mode maps Hi-C short reads on a Hifi assembly graph creating fully haplotype resolved assemblies containing haplotigs. Hi-C integrated Hifiasm has been applied on humans and other vertebrates [9] and plant genomes to create haplotype resolved genomes [20]. A previous study on a genome of a diploid African cassava cultivar has achieved a high accuracy and contiguity of two chromosome scale haplotypes with a low percentage of misjoined haplotigs from different chromosomes with hifiasm with Hi-C mode [20]. This reveals that the Hifiasm (Hi-C mode) works well for diploid genomes to generate haplotype resolved assemblies but requires further validation to detect haplotype specific variations with high confidence.

For citrus, haplotype resolved genomes have been generated for lemon using Falcon in combination with purge haplotig pipeline [14] and sweet orange using Falcon-unzip [15] to date. We produced two haplotypes for *C. australis* generated by Hi-C integrated mode of Hifiasm which facilitated the identification of variations in gene number and chromosomal lengths with respect to the collapsed assembly. Significant differences between haplotypes in terms of assembly lengths and annotated genes have previously been reported in other plant genomes with different assemblers [21,22]. The differences between the haplotypes could possibly be due to the actual biological variations or due to assembly artifacts which should be verified by further analysis.

Citrus is threatened by a plethora of pests and diseases, with HLB being the most destructive disease in the world [23]. Recently, a novel class of stable amicrobial peptide has been isolated and characterized from HLB resistant species *C. australasica* and *P. trifoliata* which differs mostly from other antimicrobial peptides produced by HLB susceptible species in terms of length. The short peptide having 67 aa and long peptides (109 aa) were detected in HLB resistant plants, however the susceptible species were only detected with a long aa (118 aa and 109 aa). This novel peptide containing two cystine residues and α-helix2 domain can cause cytosol leakage and cell lysis of the disease causing *Candidatus liberibacter* bacteria, thus suppress their growth and induce the immunity in host plants, preventing further bacterial infections [7].

*C. australis* has also been characterized as a resistant species by previous extensive field experiments [4]. The present gene annotation identified the gene g9664 which is homologous to the short novel peptide of *C. australasica* with two transcripts, encoding long aa residues which are not present in susceptible species. The 67 aa sequence is present within these long peptides with only a few amino acid substitutions or insertions. Therefore, it is quite possible that these larger precursor proteins in *C. australis* may later be modified by proteolytic cleavage to produce the 67 SAMP versions of resistant cultivars. It has been reported that different accessions of some species such as *P. trifoliata* can have different degrees of resistance to HLB [4] and this reveals the importance of having a complete genetic picture of all the available accessions or varieties in a species to capture all the allelic variations among them. The gene g9664 could be a potential candidate for resistance against HLB and it’s worth monitoring the expression of this gene in response to HLB infection to further validate the function of the gene.

We also annotated three other types of defence related genes which might play important roles against HLB. Leucine rich repeat containing proteins play pivotal roles providing innate immunity in plants by facilitating pathogen recognition [24]. A previous study on HLB resistance of *P. trifoliata* has identified NBS-LRR genes, a most common type of plant disease resistant genes, and a rapidly evolving gene family containing non-NBS type LRR genes which might play crucial roles in disease resistance [25]. Another study on an HLB resistant transgenic line identified enhanced expression levels of leucine-rich repeat receptor kinases (LRR-RKs) compared to susceptible plants revealing the importance of these genes for HLB resistant in citrus [26]. We identified 76 genes encoding LRR proteins in *C. australis* genome which might be crucial for HLB resistance. In addition, the annotation explored 17 guanine nucleotide binding proteins which confer resistance against biotic stresses [27] and 13 pathogenesis-related proteins which have previously been identified as highly upregulated genes in HLB infected *C. australasica* plants [28]. Currently no reference genome is available for Australian limes, therefore the high-quality genome presented in this study paves the way for comparative genomics with other HLB resistant citrus species to fully understand the resistance mechanisms for HLB.

Sugar and organic acids are major attributes of fruit flavour [29]. Some citrus species including limes are highly acidic compared to the cultivated citrus species and the reduced acidity is considered to be an important trait during citrus domestication [30]. Hyper acidification of vacuolar epidermal cells of highly acidic species including limes is controlled by two interacting P-ATPase, *CitPH1* (P3B-ATPase - Mg2+ pump) and *CitPH5* (P3A-ATPase - H+ pump). [19]. *C. australis* is a wild lime with an edible acidic pulp. A good flavour characterized by a proper balance of sourness and sweetness is vital in improving the citrus market value [31]. Australian wild limes are good genomic resources to be tested in breeding novel types of acidic fruits. We here identified the two key genes, *PH1* and *PH5* and their transcription regulators in *C. australis*. Besides the two key genes, nine other genes encoding P-type H(+)-exporting transporters reside in Chr 4 and 6 which might be the other vacuolar H+-ATPases (V-ATPase) driving the citrate import into vacuoles through H+ gradient. A few amino acid substitutions of PH1 protein identified in this study may be involved in the acidification of vacuoles in *C. australis* and may differentiate the acidic and sweet citrus species. The other sweet and acidic citrus species are required to validate these amino acid sequence variations with confidence; however, this is limited due to the lack of high-quality genomes for wild acidic citrus species. Aconitase (CitAco) would be a good target for developing reduced acidic cultivars through gene editing. The genetic loci identified in this study will provide valuable molecular markers for marker assisted breeding for selecting good flavours in *C. australis*. The annotated gene sequences will also be a good resource to understand the sequence variations of these genes in wild limes with compared to the cultivated species.

Leaf oils have previously been characterized in Australian wild limes where α-pinene is the dominant compound in *C. australis* (68-79%) [2]. So far, none of the genes specific to the generation of terpenes have been identified and characterized in *C. australis*. Here we have identified and functionally characterized the genes encoding terpene synthesis genes scattered across the *C. australis* genome. We couldn’t identify the specific genes for α-pinene in *C. australis* genome, however, TPS27, probable terpene synthase 6 or probable terpene synthase 9 could be potential genes encoding α-pinene. In addition to the principal component α-pinene, other monoterpenes including β-pinene, myrcene, limonene, β-phellandrene, linalool and sesquiterpenes such as bicyclogermacrene, globulol, and viridiflorol have been isolated from *C. australis* leaves previously [2]. Limonin possess anticarcinogenic and anti-HIV activities [32] while α-pinene possess numerous benefits including neuroprotective effects [33], and anticancer activities [34]. Identification and functional characterization of genes for specific volatile compounds would provide breeders with direction for developing cultivars with increased levels of beneficial compounds that would in turn improve the consumer demand.

The high-quality *C. australis* genome presented here provides a good resource to improve citrus quality attributed traits through genetic breeding. This genome provides unprecedented opportunities for comparative genomics with other Australian wild limes and commercial citrus species to further understand the species-specific traits, mechanisms underlining biotic stresses, and their evolutionary relationships which will support the applied breeding efforts. The availability of good reference genomes for Australian native citrus species further facilitates the assembly of pangenomes to explore the existing genomic diversity among the species.

## MATERIALS AND METHODS

### Sample collection, DNA and RNA extraction and sequencing, Hi-C sequencing and Flow cytometry

Young fresh leaves of *Citrus australis* were collected from a plant grown in a glasshouse at The University of Queensland, Australia (−27.495859, 153.010139) which was sourced from Ross Evans Nursery in Kenmore, QLD, 4069. Total genomic DNA was extracted from pulverized leaf tissues using a CTAB (Cetyltrimethyl ammonium bromide) DNA extraction protocol [35]. PacBio Sequencing was performed on two PacBio Sequel II SMRT cells using the circular consensus sequence method to generate HiFi reads at The Australian Genome Research Facility (AGRF), University of Queensland. Total RNA was extracted from leaves using Trizol and Qiagen kit methods [36]. RNA was sequenced at the AGRF, The University of Queensland, Australia. Fresh young leaves were collected from the same plant for Hi-C sequencing and Flow cytometry. Hi-C sequencing was performed at The Ramaciotti Centre for Genomics, University of New South Wales, Australia. The Hi-C library preparation and analysis were done using Phase Genomics Proximo Plant Hi-C version 4.0. Flow cytometry was performed at the University of Queensland using the BD Biosciences LSR II Flow Cytometer and analysed with the FlowJo software package. Briefly, fresh C. australis was co-chopped with the reference standard, Macadamia tetraphylla (presumed size 796 Mb) in Arumuganathan and Earle buffer (https://doi.org/10.1007/BF02672073). Nuclei were gently filtered through a pre-soaked 40-µm nylon mesh and stained with 50µg/mL of propidium iodide and 50µg/ml of RNase A. Three biological replicates were performed on three different days.

### Genome assembly

PacBio high fidelity (HiFi) reads were assembled using the Hifiasm Denovo assembler [9] in three modes to produce contigs. In the first mode, Hifi reads were used alone with built-in duplication parameters. In the second mode, Hifi reads were used with --primary option in Hifiasm. In the third mode, Hifi reads were used with Hi-C reads using Hi-C partition options in hifiasm. The contig assemblies were scaffolded using SALSA tool [37]. Genome assembly completeness was assessed using Benchmarking Universal Single-Copy Orthologs against 425 single copy orthologs in viridiplantae lineage (BUSCO v5.2.2) [38] and the contiguity was assessed using QUAST (version 5.0.2) [39]. The longest nine scaffolds were assigned to chromosomes that correspond with previously published chromosome-scale genomes; *C. sinensis* v.03 [40], *C. maxima* [41] and *C. limon* [14]. The synteny among the genomes were visualized with D-Genies v.1.4 [42]. The pseudochromosomes were characterized in terms of telomere repeats [43] and Ribosomal RNA gene repeats [44]. Chloroplast genome which was assembled by GetOrganelle toolkit v.1.7.5 [45] was compared with three sets of scaffolds in dotplots using Dgenies to identify the proportion of the nuclear genome covered by the chloroplast genomic fragments. The K-mer analysis was performed using Jellyfish (v2.2.10) [46] and the histograms were further analyzed using genomescope web tool (http://qb.cshl.edu/genomescope/) [17] with the maximum k-mer coverage set to 1000,000. The genome size was also estimated using kmergenie with K=97 [47] and Flow cytometry [48].

### Genome annotation

Repeat elements were de novo detected by Repeatmodeler2 version 2.0.1 [49] and masked by Repeatmasker version 4.0.9_p2 [18]. Repeat masking was performed in three options: soft masking, hard masking, and hard masking with “-nolow” option. Quality and adapter trimmed RNA-seq reads were aligned to the unmasked and masked genomes using HISAT2 [50]. Evidence based gene prediction was performed by Braker v2.1.6 [51]. BUSCO was used to assess the genome annotation completeness.

Functional annotation was performed in OmicsBox 2.2.4. CDS sequences were subjected to BLASTX program with viridiplantae taxonomy against the Non-redundant protein sequences database with an e-value of 1.0E-10. CDS were ran through InterProScan and GO terms were retrieved for all the hits obtained by BLAST search using Gene Ontology mapping with Blast2GO annotation. InterProScan and Blast2go annotations were then combined, and the GO terms retrieved from InterProScan were merged with those retrieved from Blast2go annotations. CDS sequences with no BLAST hits obtained from Genemark trained Augustus were extracted and run through coding potential assessment using the prebuilt model of *Arabidopsis thaliana* and by creating citrus specific models to distinguish the transcripts that were coding and non-coding.

### Identification of citrus specific genes

Genes involved in the production of antimicrobial peptides, defense, volatile compounds and acidity regulation were identified by BLAST homology search with an e-value of 1.0E-10, in CLC (Qiagen, USA). The homologs of *C. australis* and other citrus species were compared by sequence alignment using Clone Manager Ver. 9. The peptide sequences from citrus and other species were obtained from NCBI, citrus genome database (CGD) and Citrus Pan-genome to Breeding Database (CPBD) [52].

## Acknowledgements

This project was funded by the Hort Frontiers Advanced Production Systems Fund as part of the Hort Frontiers strategic partnership initiative developed by Hort Innovation, with co-investment from The University of Queensland, and contributions from the Australian Government and Bioplatforms Australia. UN was supported by a graduate scholarship from The University of Queensland. The authors acknowledge The University of Queensland Research Computing Centre (UQ-RCC) for providing all the computing resources for the study and the Flow Cytometry Facility at the Queensland Brain Institute.

## Author contributions

RH, AF, AKM supervised, managed the project, advised and supported data analysis, and data interpretation. AF advised on experiments and AKM supported the genome assembly work. UN and PM performed DNA extractions. UN conducted RNA extractions, data analysis and data interpretation. LC contributed flow cytometry analysis. The manuscript was organized and written by UN. LC contributed to the manuscript with data interpretation of flow cytometry analysis. All authors approved the submitted version.

## Data availability statement

Raw sequence data generated in this study have been deposited in NCBI Sequence Read Archive (SRA) under BioProject PRJNA910964 and BioSample SAMN32155198 with an accession ID of SRR22742835 for RNA-seq and SRR22793114 for whole genome short read data. The whole genome sequence data reported in this paper have been deposited in the Genome Warehouse in National Genomics Data Center [53,54], Beijing Institute of Genomics, Chinese Academy of Sciences / China National Center for Bioinformation, under accession number GWHBQDX00000000, BioProject [PRJCA013889], and Biosample [SAMC1020632] that is publicly accessible at https://ngdc.cncb.ac.cn/gwh.

