## Supplemental material for "Haplotype resolved chromosome level genome assembly of *Citrus australis* reveals disease resistance and other citrus specific genes"

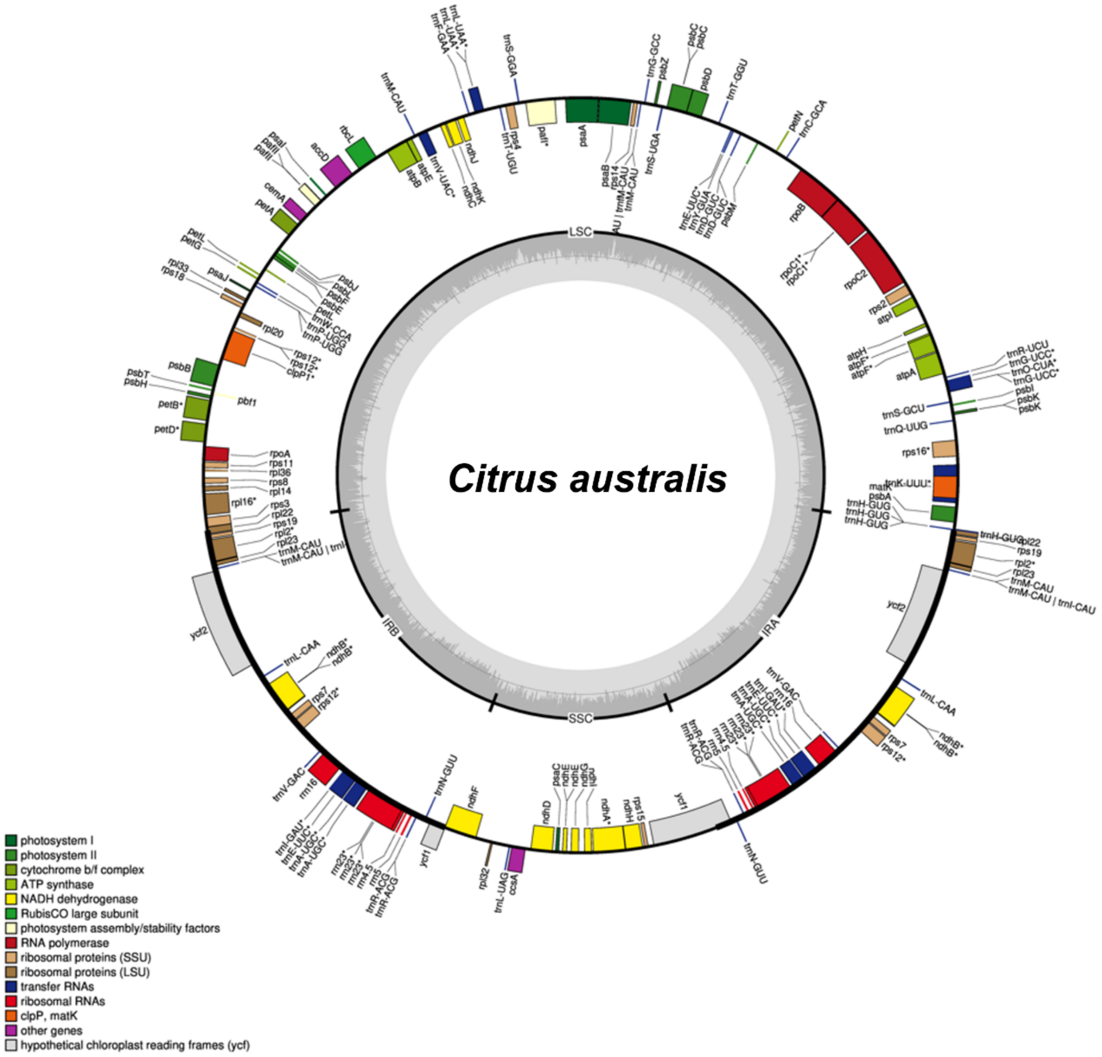

**Supplementary Fig. S1** Chloroplast genome map of *C. australis*. *C. australis* showed the typical quadripartite structure of the chloroplast genome. The genes belonging to different functional groups are shown in different colors. The thicker lines indicate the extent of the IR regions separating the LSC and SSC regions. Genes inside the circle are transcribed in clockwise direction whereas the genes outside the circle are transcribed in counter-clockwise direction. LSC: Large Single-Copy, SSC: Small Single-Copy, IR: Inverted Repeat regions

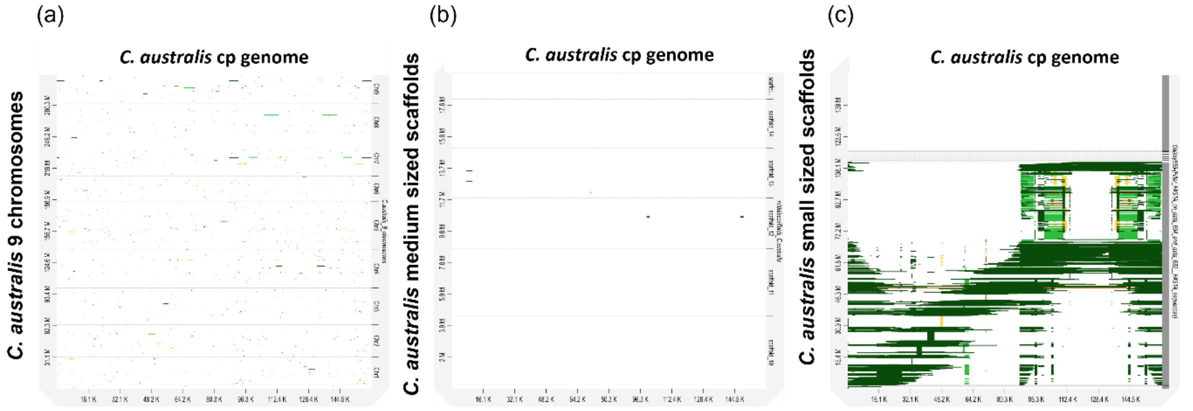

**Supplementary Fig. S2** Alignments of *C. australis* complete chloroplast genome with different sets of assembled scaffolds. Large parts of the scaffolds 27-4642 showed high sequence similarities with the chloroplast genome.

Among the medium sized scaffolds, only 12 and 13 scaffolds contain small fragments of the chloroplast genome. Sequence similarities with some parts of the top 9 scaffolds with the chloroplast genome indicate the insertion of chloroplast sequences within the nuclear genome.

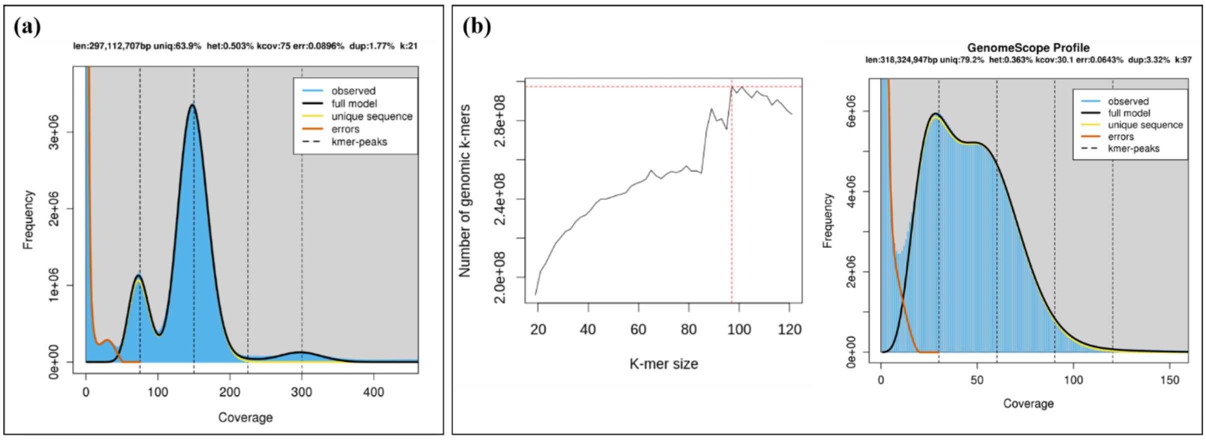

**Supplementary Fig. S3** Genome size estimation based on K-mer profile spectrum analysis (a) K-mer profile showed a distinct bimodal profile which is characteristic to diploid heterozygous genomes. The fist peak reveals heterozygous k-mers while the second peak indicates the homozygous k-mers. The relative height between the two peaks is an indication of the heterozygosity level in the genome. K=21 estimated 297 as the genome size with 0.503% heterozygosity. (b) Kmergenie predicted the best k as 97 and the genome size was estimated as 318 bp with 0.363% heterozygosity using k=97 in genomescope.

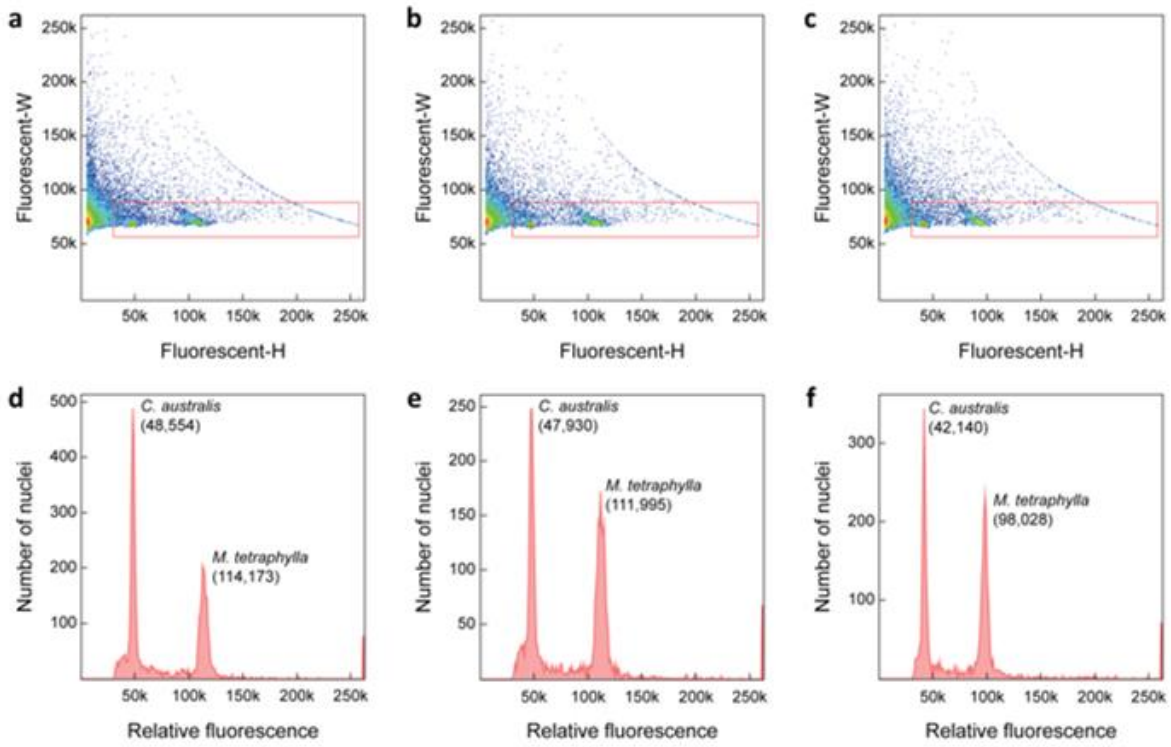

**Supplementary Fig. S4** The relative fluorescence intensities of plant nuclei isolated from *Citrus australis* co-chopped with the standard *Macadamia tetraphylla* for three biological replicates. A gating map was used to estimate debris and nuclei clumps using fluorescent height and width ratios (a-c; red box). The peak fluorescence intensity values on gated histograms were used to calculate nuclear DNA content (d-f).

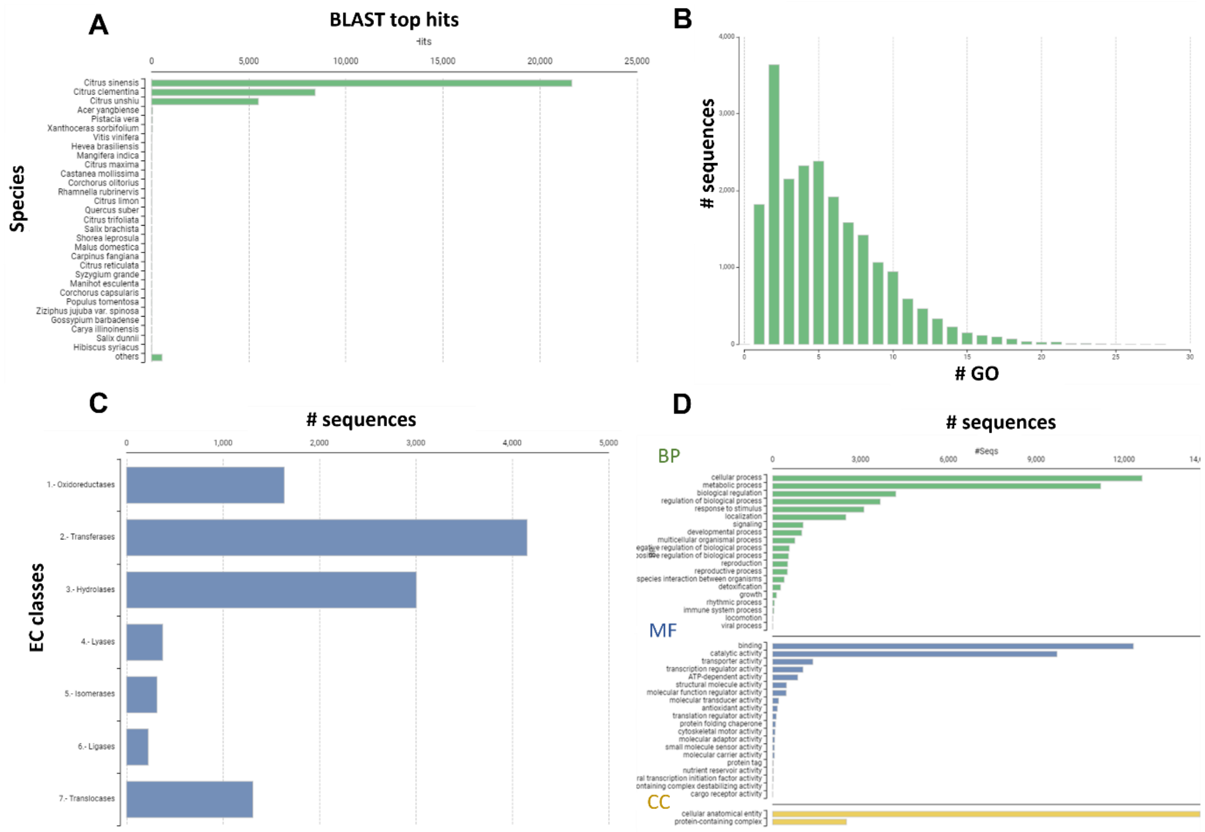

**Supplementary Fig. S5** (A) Top hit species distribution (B) GO Mapping Distribution showing the distribution of the amount of Gene Ontology candidate terms assigned to each sequence during the GO Mapping step. (C) Enzyme code distribution showing the number of sequences encoding different enzyme classes (D) GO terms for all 3 categories: Molecular function, biological function and cellular process.

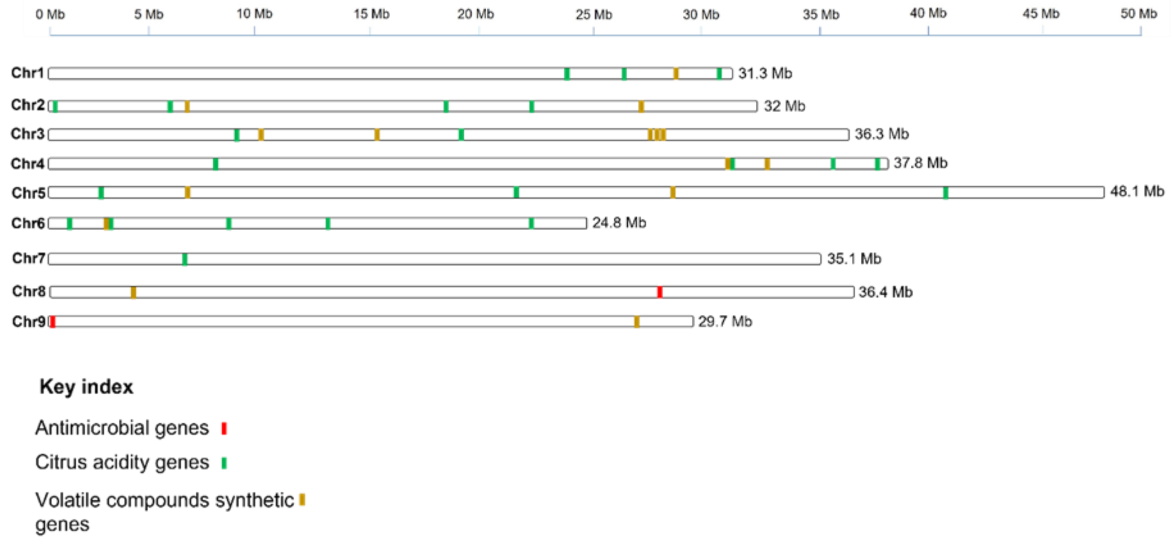

**Supplementary Fig. S6** Chromosomal positions of antimicrobial genes, acidity related genes and volatile compounds synthesis genes. Two antimicrobial genes are located in Chr 8 and 9. 25 acidity related genes are dispersed in all the 9 chromosomes and 26 volatile compound synthesis genes are present in all except Chr7.

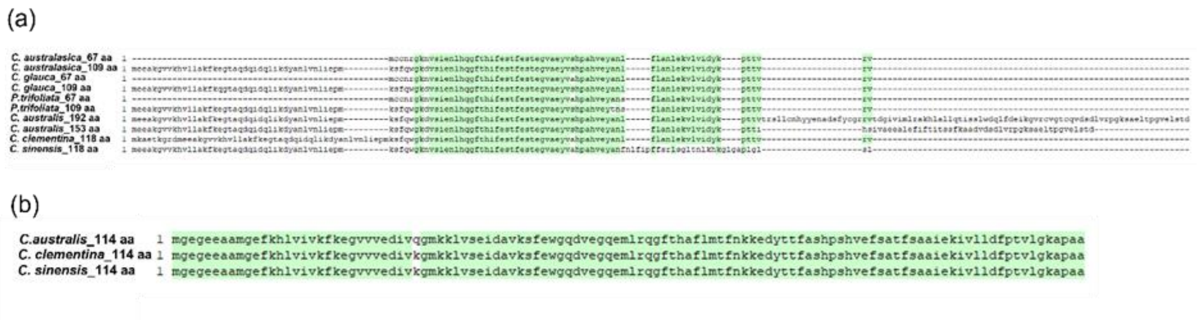

**Supplementary Fig. S7** Antimicrobial proteins exist in *C. australis* genome. (a) g9664 gene resides in chromosome 9 and encodes two transcripts giving rise to two stress-response A/B barrel domain-containing protein HS1. One protein is 153 aa lengthy and has 37% sequence similarity with 67 SAMPs found in HLB resistant species (*C. australasica*, *C. glauca* and *P. trifoliata*). The other transcript encodes 192 aa protein which has 31% sequence similarity with 67 SAMPs. These antimicrobial proteins are different from those of HLB susceptible species (*C. sinensis* and *C. clementina*). (b) g2059 gene resides in chromosome 8 encodes 114 aa peptide which is identical to those produced by HLB susceptible species (*C. sinensis* and *C. clementina*) except for one SNP at 31 aa positions.

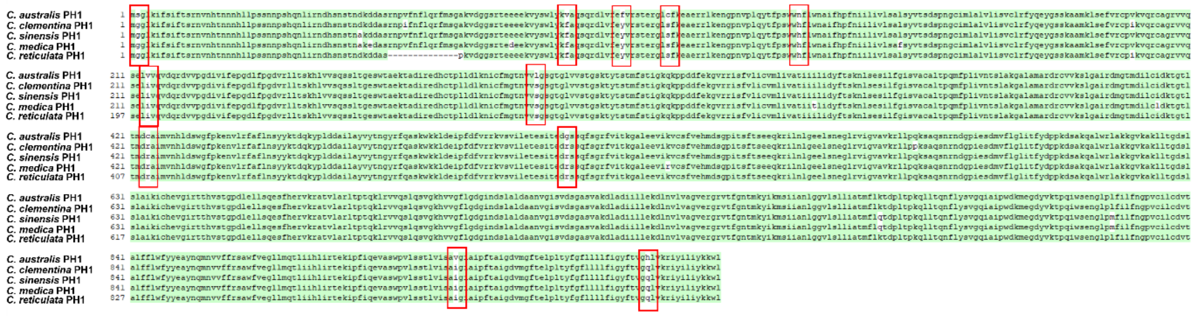

**Supplementary Fig. S8** Alignment of PH1 protein of *C. australis* with other cultivated citrus species. Red color boxes show amino acid substitutions among *C. australis* (acidic taste) and other cultivated citrus species (sweet taste) in PH1 protein.

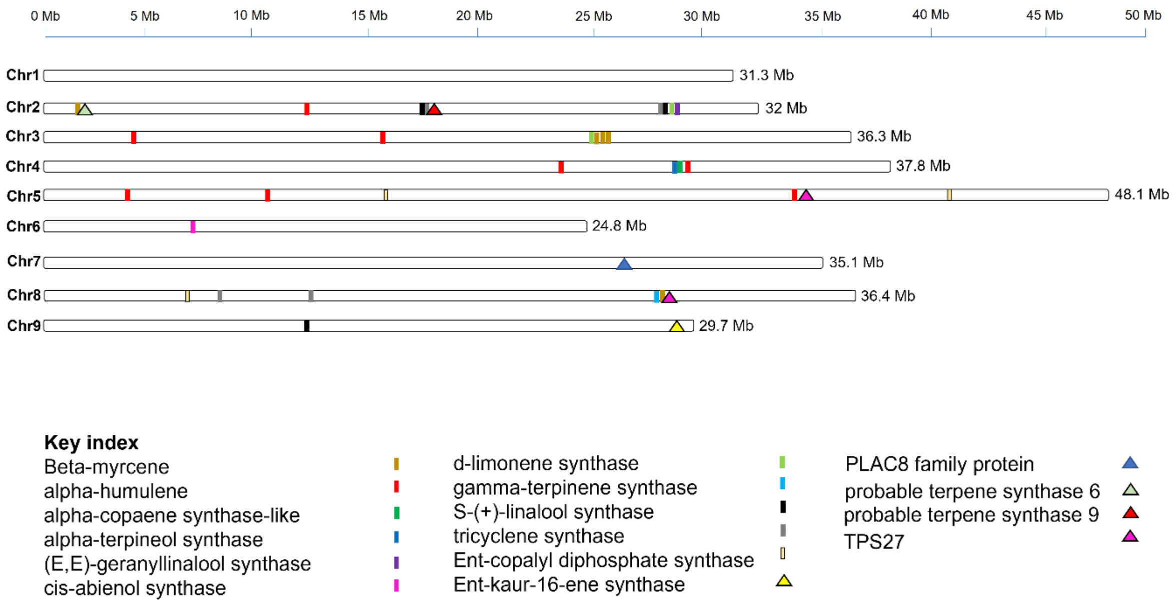

**Supplementary Fig. S9** 79 Terpene Synthase (TPS) genes in *C. australis* genome. 37 genes produce monoterpenes, 24 genes are involved in sesquiterpenes, and nine genes are involved in the synthesis of diterpenoids

**Supplementary Table S1** Summary of sequence data used for genome assembly

| Platform | PacBio (HiFi) | | Illumina | HiC | RNA-seq |
|---|---|---|---|---|---|
| | **SMRT cell 1** | **SMRT cell 2** | | | |
| Number of reads | 2.21 M | 2.05 M | 481 M | 656 M | 250 M |
| Yield (bp) | 30.7 Gb | 27.8 Gb | 67.2 Gb | 99.1 Gb | 37.6 Gb |
| Read quality (median) | Q32 | Q32 | - | | |
| coverage | 90.17 | 81.83 | 177 | 292 | 110 |

**Supplementary Table S2** Contiguity and completeness of three *C. australis* scaffold level assemblies generated by Hi-C data

| | | | | Assembly contiguity | | | | Assembly completeness (%) | | | | |
|---|---|---|---|---|---|---|---|---|---|---|---|---|
| **Options in Hifiasm** | **Type of reads used** | **Type of assembly** | **Number of scaffolds** | **Total length (Mb)** | **Largest contig (Mb)** | **N50 (Mb)** | **L50** | **Complete BUSCOs** | **Complete & single-copy BUSCOS** | **Complete & duplicated BUSCOS** | **Fragmented BUSCOS** | **Missing BUSCOS** |
| 1. Hifi reads in default | Hifi reads | Collapsed | 4663 | 485 | 48 | 29.7 | 7 | 98.8 | 98.6 | 0.2 | 0.5 | 0.7 |
| 2. Hifi reads with primary option | Hifi reads | Primary | 4618 | 487 | 48 | 31.3 | 7 | 98.8 | 98.6 | 0.2 | 0.5 | 0.7 |
| 3. Hi-C integrated assembly | Hifi reads + Hi-C reads | Collapsed | 4642 | 486 | 48 | 31.3 | 7 | 98.8 | 98.6 | 0.2 | 0.5 | 0.7 |
| | | hap1 | 4393 | 470 | 47 | 30 | 7 | 98.8 | 98.6 | 0.2 | 0.5 | 0.7 |
| | | hap2 | 1476 | 357 | 47 | 30.6 | 5 | 97.4 | 97.2 | 0.2 | 0.5 | 2.1 |

**Supplementary Table S3** Characteristics of nine chromosome scale pseudomolecules

| Chromosome number | Size (Mb) | Corresponding contig | Terminal sequence 1 | Terminal sequence 2 |
|---|---|---|---|---|
| 1 | 31.3 | ptg 7 | Telomere | Telomere |
| 2 | 32 | ptg 5 | - | Telomere |
| 3 | 36.3 | ptg 4 | Telomere | Telomere |
| 4 | 37.8 | ptg 13 | - | Telomere |
| 5 | 48.1 | ptg 3 | Telomere | Telomere |
| 6 | 24.8 | ptg 9 + ptg 48 | Telomere | - |
| 7 | 35.1 | ptg 6 | Telomere | Telomere/5s rRNA |
| 8 | 36.4 | ptg 11 + ptg 30 | Telomere | - |
| 9 | 29.7 | ptg 1 | Telomere | - |

**Supplementary Table S4** Number of regions occupied by different repeat regions in individual chromosomes

| Repeat class | Chr1 | Chr2 | Chr3 | Chr4 | Chr5 | Chr6 | Chr7 | Chr8 | Chr9 | Total |
|---|---|---|---|---|---|---|---|---|---|---|
| LTR copia | 2,352 | 2,340 | 2,869 | 2,455 | 3,799 | 2,022 | 2,154 | 2,628 | 2,836 | 23,455 |
| LTR caulimovirus | 679 | 469 | 901 | 722 | 616 | 826 | 691 | 675 | 699 | 6,278 |
| LTR ERV1 | 28 | 30 | 52 | 34 | 55 | 35 | 48 | 60 | 32 | 374 |
| LTR gypsy | 2,040 | 2,035 | 2,390 | 2,707 | 2,959 | 1,822 | 2,102 | 2,698 | 2,439 | 21,192 |
| LTR Ngaro | 40 | 39 | 57 | 45 | 102 | 23 | 36 | 41 | 48 | 431 |
| LTR_Pao | 26 | 20 | 16 | 27 | 38 | 18 | 17 | 19 | 18 | 199 |
| LTR_unknown | 1,782 | 1,620 | 2,377 | 1,656 | 2,659 | 1,326 | 1,618 | 1,864 | 2,026 | 16,928 |
| LINE/L1 | 621 | 654 | 741 | 605 | 1,083 | 513 | 666 | 643 | 679 | 6,205 |
| RC/Helitron | 361 | 143 | 1,520 | 171 | 475 | 116 | 80 | 109 | 127 | 3,102 |
| DNA/CMC-EnSpm | 371 | 382 | 401 | 278 | 623 | 302 | 286 | 409 | 434 | 3,486 |
| DNA/CMC-Transib | 5 | 1 | 8 | 0 | 213 | 0 | 0 | 2 | 0 | 229 |
| DNA/hAT-Ac | 639 | 646 | 660 | 726 | 919 | 549 | 675 | 759 | 821 | 6,394 |
| DNA/hAT-Tag1 | 53 | 61 | 84 | 61 | 117 | 60 | 76 | 54 | 43 | 609 |
| DNA/hAT-Tip100 | 148 | 194 | 246 | 160 | 204 | 136 | 164 | 179 | 168 | 1,599 |
| DNA/Maverick | 171 | 183 | 214 | 154 | 251 | 147 | 197 | 196 | 179 | 1,692 |
| DNA/Merlin | 90 | 158 | 84 | 118 | 182 | 98 | 98 | 115 | 54 | 997 |
| DNA/MULE-MuDR | 583 | 655 | 540 | 626 | 994 | 468 | 568 | 639 | 592 | 5,665 |
| DNA/PIF-Harbinger | 110 | 142 | 179 | 128 | 173 | 108 | 99 | 113 | 106 | 1,158 |
| Unknown/helitron | 5 | 7 | 7 | 1 | 252 | 0 | 1 | 17 | 13 | 303 |
| Unknown | 21,062 | 20,750 | 26,399 | 19,085 | 33,793 | 15,770 | 18,046 | 19,863 | 20,563 | 195,331 |
| Low complexity | 1,608 | 1,867 | 1,841 | 1,517 | 2,552 | 1,294 | 1,492 | 1,390 | 1,428 | 14,989 |
| Satellite | 15 | 26 | 17 | 12 | 31 | 11 | 8 | 10 | 22 | 152 |
| Simple repeats | 7,884 | 9,305 | 9,156 | 7,859 | 12,990 | 6,506 | 7,650 | 6,972 | 7,205 | 75,527 |
| snRNA | 45 | 30 | 26 | 41 | 55 | 24 | 38 | 25 | 47 | 331 |
| tRNA | 7 | 14 | 9 | 5 | 7 | 4 | 4 | 5 | 1 | 56 |
| rRNA | 55 | 57 | 53 | 53 | 64 | 955 | 45 | 58 | 56 | 1,396 |

**Supplementary Table S5** Length (bp) associated with repetitive regions

| **Repeat class** | **Chr1** | **Chr2** | **Chr3** | **Chr4** | **Chr5** | **Chr6** | **Chr7** | **Chr8** | **Chr9** | **Total** |
|---|---|---|---|---|---|---|---|---|---|---|
| LTR copia | 2,336,812 | 2,289,113 | 3,076,061 | 2,581,927 | 3,719,867 | 2,093,940 | 2,241,718 | 2,773,021 | 2,885,230 | 23,997,689 |
| LTR caulimovirus | 784,865 | 576,416 | 982,151 | 853,410 | 715,525 | 732,223 | 742,829 | 774,210 | 811,827 | 6,973,456 |
| LTR ERV1 | 5,453 | 6,788 | 11,100 | 6,645 | 11,807 | 6,719 | 9,419 | 18,204 | 5,809 | 81,944 |
| LTR gypsy | 2,907,587 | 3,146,954 | 3,459,039 | 3,943,944 | 4,200,606 | 2,574,709 | 3,078,440 | 3,903,342 | 3,316,240 | 30,530,861 |
| LTR Ngaro | 9,370 | 9,982 | 12,819 | 9,370 | 20,472 | 5,386 | 8,828 | 9,797 | 9,484 | 95,508 |
| LTR_Pao | 4,670 | 4,085 | 3,967 | 5,305 | 6,985 | 3,466 | 3,667 | 4,631 | 4,931 | 41,707 |
| LTR_unknown | 617,823 | 532,202 | 829,884 | 596,032 | 908,216 | 493,499 | 534,511 | 645,519 | 696,951 | 5,854,637 |
| LINE/L1 | 543,950 | 560,770 | 639,525 | 547,405 | 934,110 | 422,630 | 519,239 | 504,295 | 598,074 | 5,269,998 |
| RC/Helitron | 205,382 | 65,111 | 734,362 | 77,616 | 316,610 | 54,586 | 29,489 | 64,791 | 45,017 | 1,592,964 |
| DNA/CMC-EnSpm | 171,997 | 153,035 | 182,263 | 132,540 | 327,522 | 159,805 | 137,235 | 231,641 | 256,460 | 1,752,498 |
| DNA/CMC-Transib | 6,406 | 42 | 3,256 | 0 | 74,625 | 0 | 0 | 661 | 0 | 84,990 |
| DNA/hAT-Ac | 221,898 | 243,219 | 238,076 | 286,012 | 335,020 | 219,508 | 282,292 | 313,983 | 381,211 | 2,521,219 |
| DNA/hAT-Tag1 | 8,153 | 10,946 | 11,365 | 8,491 | 16,763 | 8,959 | 11,381 | 7,770 | 5,932 | 89,760 |
| DNA/hAT-Tip100 | 55,357 | 74,840 | 92,171 | 58,221 | 76,193 | 46,053 | 60,155 | 76,756 | 64,320 | 604,066 |
| DNA/Maverick | 58,081 | 63,886 | 70,232 | 48,149 | 80,735 | 51,739 | 65,061 | 80,120 | 62,872 | 580,875 |
| DNA/Merlin | 52,962 | 88,313 | 52,990 | 74,233 | 110,928 | 57,152 | 57,759 | 63,294 | 30,791 | 588,422 |
| DNA/MULE-MuDR | 380,918 | 464,764 | 377,672 | 415,144 | 723,589 | 302,019 | 412,970 | 521,269 | 454,163 | 4,052,508 |
| DNA/PIF-Harbinger | 35,392 | 55,236 | 52,484 | 48,694 | 60,597 | 54,241 | 34,569 | 29,509 | 46,152 | 416,874 |
| Unknown/helitron | 1,636 | 2,034 | 2,259 | 308 | 78,134 | 0 | 292 | 5,919 | 4,600 | 95,182 |
| Unknown | 7,677,973 | 6,066,827 | 8,350,956 | 13,525,397 | 10,651,958 | 4,514,240 | 12,192,940 | 12,901,151 | 6,276,808 | 82,158,250 |
| Low complexity | 77,734 | 83,867 | 85,114 | 69,213 | 116,276 | 58,612 | 66,982 | 63,814 | 65,129 | 686,741 |
| Satellite | 1,931 | 2,848 | 1,416 | 1,201 | 3,853 | 931 | 591 | 1,085 | 2,603 | 16,459 |
| Simple repeats | 312,433 | 341,035 | 361,998 | 296,169 | 510,777 | 242,600 | 297,209 | 267,519 | 288,772 | 2,918,512 |
| snRNA | 9,917 | 7,128 | 5,513 | 9,213 | 9,193 | 4,376 | 8,121 | 4,686 | 8,470 | 66,617 |
| tRNA | 593 | 1,271 | 794 | 428 | 640 | 297 | 326 | 392 | 82 | 4,823 |
| rRNA | 14,462 | 15,048 | 32,676 | 17,637 | 23,717 | 108,758 | 15,108 | 17,619 | 18,513 | 263,538 |

**Supplementary Table S6** Genes predicted with quality trimmed only and quality and adapter trimmed RNA-seq evidence

| Trimming options for RNA-seq data | | Genemark trained Augustus | | Genemark only | | Braker (Augustus + Genemark) | |
|---|---|---|---|---|---|---|---|
| | Masking option | Number of genes | Number of CDS | Number of genes | Number of CDS | Number of genes | Number of CDS |
| Quality trimmed | No masked | 67223 | 70106 | 2548 | 2548 | 69,771 | 72,654 |
| | Soft masked | 29193 | 31164 | 2322 | 2322 | 31,515 | 33,486 |
| | Hard masked | 23536 | 25556 | 2059 | 2059 | 25,595 | 27,615 |
| | Hard masked_nolow | 24430 | 26554 | 2374 | 2374 | 26,804 | 28,928 |
| Quality and adapter trimmed | No masked | 69,057 | 72,815 | 2,810 | 2,810 | 71,867 | 75,625 |
| | Soft masked | 29,464 | 32,009 | 2,670 | 2,670 | 32,151 | 34,700 |
| | Hard masked | 23,775 | 26,275 | 2,198 | 2,198 | 25,973 | 28,473 |
| | Hard masked_nolow | 24,268 | 26,691 | 2,541 | 2,541 | 26,809 | 29232 |

**Supplementary Table S7** 17 Guanine nucleotide-binding proteins encoded by *C. australis* genome

| Gene name | Description | Length | GO IDs | GO names | Enzyme name |
|---|---|---|---|---|---|
| g10226 | Guanine nucleotide-binding protein subunit gamma 2 | 321 | P:GO:0007186 | P:G protein-coupled receptor signalling pathway | - |
| g10687 | extra-large guanine nucleotide-binding protein 3-like | 570 | P:GO:0007186; F:GO:0003924; F:GO:0019001; F:GO:0031683; F:GO:0046872 | P:G protein-coupled receptor signalling pathway; F:GTPase activity; F:guanyl nucleotide binding; F:G-protein beta/gamma-subunit complex binding; F:metal ion binding | nucleoside-triphosphate phosphatase |
| g1272 | Extra-large guanine nucleotide-binding protein 1 | 1908 | P:GO:0007186; F:GO:0003924; F:GO:0019001; F:GO:0031683; C:GO:0016020 | P:G protein-coupled receptor signalling pathway; F:GTPase activity; F:guanyl nucleotide binding; F:G-protein beta/gamma-subunit complex binding; C:membrane | nucleoside-triphosphate phosphatase |
| g16228 | Guanine nucleotide-binding protein subunit gamma 3 | 429 | P:GO:0007186 | P:G protein-coupled receptor signalling pathway | - |
| g16844 | Extra-large guanine nucleotide-binding protein 1 | 2739 | P:GO:0007188; F:GO:0001664; F:GO:0003924; F:GO:0005525; F:GO:0031683; C:GO:0005634; C:GO:0005834 | P:adenylate cyclase-modulating G protein-coupled receptor signalling pathway; F:G protein-coupled receptor binding; F:GTPase activity; F:GTP binding; F:G-protein beta/gamma-subunit complex binding; C:nucleus; C:heterotrimeric G-protein complex | nucleoside-triphosphate phosphatase |
| g18988 | protein phosphatase 2C and cyclic nucleotide-binding/kinase domain-containing protein | 3312 | P:GO:0007165; P:GO:0018105; F:GO:0004691; F:GO:0005524; F:GO:0017018; F:GO:0046872; C:GO:0005952 | P:signal transduction; P:peptidyl-serine phosphorylation; F:cAMP-dependent protein kinase activity; F:ATP binding; F:myosin phosphatase activity; F:metal ion binding; C:cAMP-dependent protein kinase complex | protein-serine/threonine phosphatase; cAMP-dependent protein kinase |
| g19324 | guanine nucleotide-binding protein subunit beta-like protein | 984 | F:GO:0005515; F:GO:0043022; C:GO:0015935 | F:protein binding; F:ribosome binding; C:small ribosomal subunit | - |
| g20211 | Guanine nucleotide-binding protein subunit gamma 3 | 690 | P:GO:0009737; C:GO:0005886 | P:response to abscisic acid; C:plasma membrane | - |
| g20311 | Guanine nucleotide-binding protein alpha-1 subunit | 1206 | P:GO:0007188; F:GO:0001664; F:GO:0003924; | P:adenylate cyclase-modulating G protein-coupled receptor signalling pathway; F:G protein-coupled receptor binding; F:GTPase activity; F:GTP | nucleoside-triphosphate phosphatase |

| | | | | | |
|---|---|---|---|---|---|
| | | | F:GO:0005525; F:GO:0031683; F:GO:0046872; C:GO:0005834 | binding; F:G-protein beta/gamma-subunit complex binding; F:metal ion binding; C:heterotrimeric G-protein complex | |
| g27397 | Guanine nucleotide-binding protein-like NSN1 | 1791 | F:GO:0005525; C:GO:0005730 | F:GTP binding; C:nucleolus | - |
| g27632 | guanine nucleotide-binding protein subunit gamma 1 | 324 | P:GO:0007186; P:GO:0072488; F:GO:0008519; C:GO:0005886; C:GO:0016020 | P:G protein-coupled receptor signalling pathway; P:ammonium transmembrane transport; F:ammonium transmembrane transporter activity; C:plasma membrane; C:membrane | Translocases |
| g28391 | Cyclic nucleotide-binding domain-containing protein | 2388 | P:GO:0006637; P:GO:0009062; F:GO:0005515; F:GO:0047617; C:GO:0005829 | P:acyl-CoA metabolic process; P:fatty acid catabolic process; F:protein binding; F:acyl-CoA hydrolase activity; C:cytosol | acyl-CoA hydrolase |
| g5196 | Extra-large guanine nucleotide-binding protein 3 | 2553 | P:GO:0007186; F:GO:0003924; F:GO:0019001; F:GO:0031683; F:GO:0046872 | P:G protein-coupled receptor signalling pathway; F:GTPase activity; F:guanyl nucleotide binding; F:G-protein beta/gamma-subunit complex binding; F:metal ion binding | nucleoside-triphosphate phosphatase |
| g843 | Extra-large guanine nucleotide-binding protein 1 | 3228 | P:GO:0006468; P:GO:0007186; F:GO:0003924; F:GO:0004674; F:GO:0005524; F:GO:0019001; F:GO:0031683; F:GO:0046872; C:GO:0016020 | P:protein phosphorylation; P:G protein-coupled receptor signalling pathway; F:GTPase activity; F:protein serine/threonine kinase activity; F:ATP binding; F:guanyl nucleotide binding; F:G-protein beta/gamma-subunit complex binding; F:metal ion binding; C:membrane | Transferring phosphorus-containing groups; nucleoside-triphosphate phosphatase |
| g890 | Extra-large guanine nucleotide-binding protein 1 | 3432 | P:GO:0007186; F:GO:0003924; F:GO:0019001; F:GO:0031683; F:GO:0046872 | P:G protein-coupled receptor signalling pathway; F:GTPase activity; F:guanyl nucleotide binding; F:G-protein beta/gamma-subunit complex binding; F:metal ion binding | nucleoside-triphosphate phosphatase |
| g892 | Extra-large guanine nucleotide-binding protein 1 | 3477 | P:GO:0007186; F:GO:0003924; F:GO:0019001; | P:G protein-coupled receptor signalling pathway; F:GTPase activity; F:guanyl nucleotide binding; F:G-protein beta/gamma-subunit complex binding; F:metal ion binding | nucleoside-triphosphate phosphatase |

| | | | | | |
|---|---|---|---|---|---|
| g947 | guanine nucleotide-binding protein subunit beta-2 | 948 | F:GO:0031683; F:GO:0046872 P:GO:0006950; P:GO:0007186; P:GO:0009605; P:GO:0009725; P:GO:0009791; P:GO:0048364; P:GO:0071310; F:GO:0005515; F:GO:0030159; C:GO:0005737; C:GO:0005834; C:GO:0043231 | P:response to stress; P:G protein-coupled receptor signalling pathway; P:response to external stimulus; P:response to hormone; P:post-embryonic development; P:root development; P:cellular response to organic substance; F:protein binding; F:signaling receptor complex adaptor activity; C:cytoplasm; C:heterotrimeric G-protein complex; C:intracellular membrane-bounded organelle | - |

**Supplementary Table S8** 13 pathogenesis-related proteins encoded by *C. australis* genome

| Gene name | Description | Length | GO IDs | GO names |
|---|---|---|---|---|
| g15060 | pathogenesis-related genes transcriptional activator PTI6-like | 795 | P:GO:0006355; F:GO:0003677; F:GO:0003700; C:GO:0005634 | P:regulation of DNA-templated transcription; F:DNA binding; F:DNA-binding transcription factor activity; C:nucleus |
| g16463 | pathogenesis-related protein 1-like | 480 | P:GO:0009607; C:GO:0005615 | P:response to biotic stimulus; C:extracellular space |
| g1794 | pathogenesis-related protein PR-4-like | 429 | P:GO:0042742; P:GO:0050832; P:GO:0090501; F:GO:0004540; F:GO:0008061 | P:defense response to bacterium; P:defense response to fungus; P:RNA phosphodiester bond hydrolysis; F:ribonuclease activity; F:chitin binding |
| g1797 | pathogenesis-related protein PR-4A | 432 | P:GO:0042742; P:GO:0050832; P:GO:0090501; F:GO:0004540 | P:defense response to bacterium; P:defense response to fungus; P:RNA phosphodiester bond hydrolysis; F:ribonuclease activity |
| g20662 | pathogenesis-related protein PR-1-like | 501 | P:GO:0009607; C:GO:0005615 | P:response to biotic stimulus; C:extracellular space |
| g4142 | pathogenesis-related protein 5 | 429 | P:GO:0006952; C:GO:0016020 | P:defense response; C:membrane |
| g513 | pathogenesis-related protein 1 | 480 | C:GO:0005576 | C:extracellular region |
| g520 | basic form of pathogenesis-related protein 1-like | 480 | C:GO:0005576 | C:extracellular region |
| g521 | pathogenesis-related protein 1-like | 492 | P:GO:0009607; C:GO:0005576 | P:response to biotic stimulus; C:extracellular region |
| g523 | pathogenesis-related leaf protein 6-like | 453 | C:GO:0005615 | C:extracellular space |
| g524 | pathogenesis-related protein 1 | 480 | C:GO:0005576 | C:extracellular region |
| g6255 | pathogenesis-related thaumatin-like protein 3.5 | 711 | P:GO:0006952 | P:defense response |
| g9804 | pathogenesis-related protein STH-2-like | 336 | P:GO:0006952; P:GO:0009738; P:GO:0043086; P:GO:0080163; F:GO:0004864; F:GO:0010427; F:GO:0038023; | P:defense response; P:abscisic acid-activated signaling pathway; P:negative regulation of catalytic activity; P:regulation of protein serine/threonine phosphatase activity; F:protein phosphatase inhibitor activity; F:abscisic acid binding; F:signaling receptor activity; C:nucleus; C:cytoplasm |

| | | | |
|---|---|---|---|
| | | | C:GO:0005634; C:GO:0005737 |

**Supplementary Table S9** 76 Leucine rich repeat (LRR) genes encoded by *C. australis* genome

| **Gene name** | **Description** | **Length** | **GO IDs** | **GO names** | **Enzyme name** |
|---|---|---|---|---|---|
| g10165 | leucine-rich repeat extensin-like protein 5 | 651 | C:GO:0005886; C:GO:0016020; C:GO:0031225 | C:plasma membrane; C:membrane; C:obsolete anchored component of membrane | |
| g10634 | putative leucine-rich repeat receptor-like protein kinase | 3450 | P:GO:0006468; F:GO:0004672; F:GO:0005515; F:GO:0005524; C:GO:0005886; C:GO:0016020 | P:protein phosphorylation; F:protein kinase activity; F:protein binding; F:ATP binding; C:plasma membrane; C:membrane | Transferring phosphorus-containing groups |
| g10961 | putatively inactive leucine-rich repeat receptor-like protein kinase | 2499 | P:GO:0006468; F:GO:0004672; F:GO:0005515; F:GO:0005524; C:GO:0005886; C:GO:0016020 | P:protein phosphorylation; F:protein kinase activity; F:protein binding; F:ATP binding; C:plasma membrane; C:membrane | Transferring phosphorus-containing groups |
| g11274 | leucine-rich repeat receptor-like tyrosine-protein kinase PXC3 | 2667 | P:GO:0006468; F:GO:0004672; F:GO:0005515; F:GO:0005524; C:GO:0005886; C:GO:0016020 | P:protein phosphorylation; F:protein kinase activity; F:protein binding; F:ATP binding; C:plasma membrane; C:membrane | Transferring phosphorus-containing groups |
| g11339 | probably inactive leucine-rich repeat receptor-like protein kinase At5g06940 | 2673 | P:GO:0006468; F:GO:0004672; F:GO:0005515; F:GO:0005524; C:GO:0005886; C:GO:0016020 | P:protein phosphorylation; F:protein kinase activity; F:protein binding; F:ATP binding; C:plasma membrane; C:membrane | Transferring phosphorus-containing groups |
| g11369 | leucine-rich repeat receptor protein kinase EMS1 | 3183 | P:GO:0006468; F:GO:0004674; F:GO:0005515; F:GO:0005524; C:GO:0016020 | P:protein phosphorylation; F:protein serine/threonine kinase activity; F:protein binding; F:ATP binding; C:membrane | Transferring phosphorus-containing groups |

| | | | | | |
|---|---|---|---|---|---|
| g11674 | leucine-rich repeat receptor protein kinase EMS1 | 975 | F:GO:0005515; F:GO:0016740 | F:protein binding; F:transferase activity | Transferases |
| g12132 | putative leucine-rich repeat receptor-like protein kinase | 288 | P:GO:0006468; F:GO:0000166; F:GO:0004672; F:GO:0005524; C:GO:0016020; C:GO:0016020 | P:protein phosphorylation; F:nucleotide binding; F:protein kinase activity; F:ATP binding; C:membrane; C:membrane | |
| g12465 | leucine-rich repeat extensin-like protein 2 | 642 | C:GO:0016020 | C:membrane | |
| g12687 | leucine-rich repeat receptor-like protein kinase PXC1 | 2013 | P:GO:0006468; P:GO:0009834; F:GO:0004672; F:GO:0005515; F:GO:0005524; C:GO:0016020 | P:protein phosphorylation; P:plant-type secondary cell wall biogenesis; F:protein kinase activity; F:protein binding; F:ATP binding; C:membrane | Transferring phosphorus-containing groups |
| g13313 | probably inactive leucine-rich repeat receptor-like protein kinase IMK2 | 2418 | P:GO:0006468; F:GO:0004672; F:GO:0005515; F:GO:0005524; C:GO:0016020 | P:protein phosphorylation; F:protein kinase activity; F:protein binding; F:ATP binding; C:membrane | Transferring phosphorus-containing groups |
| g13847 | probable leucine-rich repeat receptor-like protein kinase At1g35710 | 510 | F:GO:0005515 | F:protein binding | |
| g15237 | leucine-rich repeat receptor protein kinase EMS1 | 3705 | P:GO:0006468; F:GO:0004674; F:GO:0005515; F:GO:0005524; C:GO:0016020 | P:protein phosphorylation; F:protein serine/threonine kinase activity; F:protein binding; F:ATP binding; C:membrane | Transferring phosphorus-containing groups |
| g15240 | leucine-rich repeat receptor protein kinase EMS1 | 585 | P:GO:0006468; F:GO:0004674; F:GO:0005515; F:GO:0005524; C:GO:0005886; C:GO:0016020 | P:protein phosphorylation; F:protein serine/threonine kinase activity; F:protein binding; F:ATP binding; C:plasma membrane; C:membrane | Transferring phosphorus-containing groups |
| g15245 | leucine-rich repeat receptor protein kinase EMS1 | 336 | P:GO:0006468; F:GO:0004674; F:GO:0005524; C:GO:0016020 | P:protein phosphorylation; F:protein serine/threonine kinase activity; F:ATP binding; C:membrane | Transferring phosphorus-containing groups |

| | | | | | |
|---|---|---|---|---|---|
| g15246 | leucine-rich repeat receptor protein kinase EMS1 | 435 | P:GO:0006468; P:GO:0009755; F:GO:0004674; F:GO:0005515; F:GO:0005524; C:GO:0005886; C:GO:0016020 | P:protein phosphorylation; P:hormone-mediated signaling pathway; F:protein serine/threonine kinase activity; F:protein binding; F:ATP binding; C:plasma membrane; C:membrane | Transferring phosphorus-containing groups |
| g1530 | Inactive leucine-rich repeat receptor-like protein kinase CORYNE | 1062 | P:GO:0006468; F:GO:0004672; F:GO:0005524; C:GO:0005886; C:GO:0016020 | P:protein phosphorylation; F:protein kinase activity; F:ATP binding; C:plasma membrane; C:membrane | Transferring phosphorus-containing groups |
| g15307 | leucine-rich repeat receptor-like protein kinase PEPR1 | 3330 | P:GO:0006468; F:GO:0004672; F:GO:0005515; F:GO:0005524; C:GO:0016020 | P:protein phosphorylation; F:protein kinase activity; F:protein binding; F:ATP binding; C:membrane | Transferring phosphorus-containing groups |
| g15362 | putative leucine-rich repeat receptor-like serine/threonine-protein kinase At2g24130 | 2373 | P:GO:0006468; F:GO:0004672; F:GO:0005515; F:GO:0005524; C:GO:0016020 | P:protein phosphorylation; F:protein kinase activity; F:protein binding; F:ATP binding; C:membrane | Transferring phosphorus-containing groups |
| g15632 | probably inactive leucine-rich repeat receptor-like protein kinase At5g48380 | 738 | P:GO:0006468; F:GO:0004672; F:GO:0005524; C:GO:0016020 | P:protein phosphorylation; F:protein kinase activity; F:ATP binding; C:membrane | Transferring phosphorus-containing groups |
| g1651 | inactive leucine-rich repeat receptor-like serine/threonine-protein kinase At1g60630 | 1992 | P:GO:0006468; F:GO:0004672; F:GO:0005515; F:GO:0005524; C:GO:0016020 | P:protein phosphorylation; F:protein kinase activity; F:protein binding; F:ATP binding; C:membrane | Transferring phosphorus-containing groups |
| g16582 | leucine-rich repeat receptor-like serine/threonine-protein kinase BAM1 | 2286 | P:GO:0006468; P:GO:0009755; F:GO:0004672; F:GO:0005515; F:GO:0005524; C:GO:0005886; C:GO:0016020 | P:protein phosphorylation; P:hormone-mediated signaling pathway; F:protein kinase activity; F:protein binding; F:ATP binding; C:plasma membrane; C:membrane | Transferring phosphorus-containing groups |

| | | | | | |
|---|---|---|---|---|---|
| g16912 | Acidic leucine-rich nuclear phosphoprotein 32-related protein | 1365 | F:GO:0005515 | F:protein binding | |
| g16913 | Acidic leucine-rich nuclear phosphoprotein 32-related protein | 525 | C:GO:0016020 | C:membrane | |
| g18619 | probably inactive leucine-rich repeat receptor-like protein kinase At2g25790 isoform X1 | 2898 | P:GO:0006468; F:GO:0004672; F:GO:0005515; F:GO:0005524; F:GO:0032440; C:GO:0016020 | P:protein phosphorylation; F:protein kinase activity; F:protein binding; F:ATP binding; F:2-alkenal reductase [NAD(P)+] activity; C:membrane | Transferring phosphorus-containing groups; 2-alkenal reductase [NAD(P)(+)] |
| g18721 | probable leucine-rich repeat receptor-like protein kinase At1g68400 | 1923 | P:GO:0006468; F:GO:0004672; F:GO:0005515; F:GO:0005524; C:GO:0016020 | P:protein phosphorylation; F:protein kinase activity; F:protein binding; F:ATP binding; C:membrane | Transferring phosphorus-containing groups |
| g19057 | leucine-rich repeat extensin-like protein 4 | 1248 | F:GO:0005515 | F:protein binding | |
| g1927 | leucine-rich repeat and IQ domain-containing protein 1-related | 1419 | F:GO:0005515 | F:protein binding | |
| g1927 | leucine-rich repeat and IQ domain-containing protein 1-related | 1422 | F:GO:0005515 | F:protein binding | |
| g1933 | leucine-rich repeat and IQ domain-containing protein 1-related | 1458 | F:GO:0005515 | F:protein binding | |
| g20773 | putative leucine-rich repeat receptor-like serine/threonine-protein kinase At2g24130 | 2952 | P:GO:0006468; F:GO:0004672; F:GO:0005515; F:GO:0005524; C:GO:0016020 | P:protein phosphorylation; F:protein kinase activity; F:protein binding; F:ATP binding; C:membrane | Transferring phosphorus-containing groups |
| g20850 | leucine-rich repeat receptor-like protein kinase TDR | 2868 | P:GO:0018108; P:GO:2000604; F:GO:0004713; F:GO:0005515; F:GO:0005524; C:GO:0016020 | P:peptidyl-tyrosine phosphorylation; P:negative regulation of secondary growth; F:protein tyrosine kinase activity; F:protein binding; F:ATP binding; C:membrane | Transferring phosphorus-containing groups |

| | | | | | |
|---|---|---|---|---|---|
| g20878 | leucine-rich repeat extensin-like protein 2 | 2118 | F:GO:0005515 | F:protein binding | |
| g21306 | pollen-specific leucine-rich repeat extensin-like protein 2 | 327 | | | |
| g21307 | putative leucine-rich repeat receptor-like protein kinase | 252 | P:GO:0006468; F:GO:0004672; F:GO:0005524; C:GO:0016020 | P:protein phosphorylation; F:protein kinase activity; F:ATP binding; C:membrane | Transferring phosphorus-containing groups |
| g21330. | leucine-rich repeat receptor-like serine/threonine-protein kinase At2g14510 | 312 | P:GO:0018108; F:GO:0004714; F:GO:0005524; C:GO:0005615; C:GO:0005886; C:GO:0016020 | P:peptidyl-tyrosine phosphorylation; F:transmembrane receptor protein tyrosine kinase activity; F:ATP binding; C:extracellular space; C:plasma membrane; C:membrane | receptor protein-tyrosine kinase |
| g21429 | leucine-rich repeat protein 1 | 693 | F:GO:0005515 | F:protein binding | |
| g22136 | leucine-rich repeat extensin-like protein 4 | 1257 | | | |
| g22743 | putatively inactive leucine-rich repeat receptor-like protein kinase | 1425 | P:GO:0006468; F:GO:0004672; F:GO:0005524; C:GO:0016020 | P:protein phosphorylation; F:protein kinase activity; F:ATP binding; C:membrane | Transferring phosphorus-containing groups |
| g22744 | probably inactive leucine-rich repeat receptor-like protein kinase At5g48380 | 1263 | P:GO:0006468; F:GO:0004672; F:GO:0005524; C:GO:0016020 | P:protein phosphorylation; F:protein kinase activity; F:ATP binding; C:membrane | Transferring phosphorus-containing groups |
| g23111 | probable leucine-rich repeat receptor-like protein kinase At1g68400 | 2019 | P:GO:0006468; F:GO:0004672; F:GO:0005524; C:GO:0016020 | P:protein phosphorylation; F:protein kinase activity; F:ATP binding; C:membrane | Transferring phosphorus-containing groups |
| g2412 | leucine-rich repeat receptor-like serine/threonine-protein kinase At2g14510 | 438 | C:GO:0016020; C:GO:0016020 | C:membrane; C:membrane | |
| g2415 | putative leucine-rich repeat receptor-like serine/threonine-protein kinase At2g04300 | 489 | | | |
| g24839 | leucine-rich repeat receptor-like serine/threonine/tyrosine-protein kinase SOBIR1 | 1518 | P:GO:0006468; F:GO:0004674; F:GO:0005515; F:GO:0005524; | P:protein phosphorylation; F:protein serine/threonine kinase activity; F:protein binding; F:ATP binding; C:plasma membrane; C:membrane | Transferring phosphorus-containing groups |

| | | | | | |
|---|---|---|---|---|---|
| | | | C:GO:0005886; C:GO:0016020 | | |
| g25761 | putative leucine-rich repeat receptor-like serine/threonine-protein kinase | 2988 | P:GO:0006468; F:GO:0004674; F:GO:0005524; C:GO:0016020 | P:protein phosphorylation; F:protein serine/threonine kinase activity; F:ATP binding; C:membrane | Transferring phosphorus-containing groups |
| g25765 | putative leucine-rich repeat receptor-like serine/threonine-protein kinase | 315 | | | |
| g25766 | putative leucine-rich repeat receptor-like serine/threonine-protein kinase | 279 | P:GO:0006468; F:GO:0004674; F:GO:0005524; C:GO:0016020 | P:protein phosphorylation; F:protein serine/threonine kinase activity; F:ATP binding; C:membrane | Transferring phosphorus-containing groups |
| g25767 | putative leucine-rich repeat receptor-like serine/threonine-protein kinase | 3015 | P:GO:0006468; F:GO:0004672; F:GO:0005515; F:GO:0005524 | P:protein phosphorylation; F:protein kinase activity; F:protein binding; F:ATP binding | |
| g25768 | putative leucine-rich repeat receptor-like serine/threonine-protein kinase | 2676 | P:GO:0006468; F:GO:0004674; F:GO:0005515; F:GO:0005524; C:GO:0016020 | P:protein phosphorylation; F:protein serine/threonine kinase activity; F:protein binding; F:ATP binding; C:membrane | Transferring phosphorus-containing groups |
| g25772 | putative leucine-rich repeat receptor-like serine/threonine-protein kinase | 2157 | P:GO:0006468; F:GO:0004672; F:GO:0005524; C:GO:0016020 | P:protein phosphorylation; F:protein kinase activity; F:ATP binding; C:membrane | |
| g25773 | putative leucine-rich repeat receptor-like serine/threonine-protein kinase | 2865 | P:GO:0006468; F:GO:0004674; F:GO:0005515; F:GO:0005524; C:GO:0016020 | P:protein phosphorylation; F:protein serine/threonine kinase activity; F:protein binding; F:ATP binding; C:membrane | Transferring phosphorus-containing groups |
| g25807 | putative leucine-rich repeat receptor-like serine/threonine-protein kinase | 2604 | P:GO:0006468; F:GO:0004674; F:GO:0005515; F:GO:0005524; C:GO:0016020 | P:protein phosphorylation; F:protein serine/threonine kinase activity; F:protein binding; F:ATP binding; C:membrane | Transferring phosphorus-containing groups |
| g26780 | leucine-rich repeat receptor-like serine/threonine/tyrosine-protein kinase SOBIR1 | 270 | P:GO:0018108; F:GO:0004674; F:GO:0004714; | P:peptidyl-tyrosine phosphorylation; F:protein serine/threonine kinase activity; F:transmembrane receptor protein tyrosine | receptor protein-tyrosine kinase |

| | | | | | |
|---|---|---|---|---|---|
| | | | F:GO:0005524; C:GO:0005886; C:GO:0016020 | kinase activity; F:ATP binding; C:plasma membrane; C:membrane | |
| g26781 | leucine-rich repeat receptor-like serine/threonine/tyrosine-protein kinase SOBIR1 | 489 | P:GO:0010942; P:GO:0018108; P:GO:0031349; P:GO:0060862; F:GO:0004674; F:GO:0004714; F:GO:0005524; C:GO:0005886; C:GO:0016020 | P:positive regulation of cell death; P:peptidyl-tyrosine phosphorylation; P:positive regulation of defense response; P:negative regulation of floral organ abscission; F:protein serine/threonine kinase activity; F:transmembrane receptor protein tyrosine kinase activity; F:ATP binding; C:plasma membrane; C:membrane | receptor protein-tyrosine kinase |
| g26783 | leucine-rich repeat receptor-like serine/threonine/tyrosine-protein kinase SOBIR1 | 408 | P:GO:0018108; F:GO:0004674; F:GO:0004714; F:GO:0005524; F:GO:0017018; F:GO:0046872; C:GO:0005886 | P:peptidyl-tyrosine phosphorylation; F:protein serine/threonine kinase activity; F:transmembrane receptor protein tyrosine kinase activity; F:ATP binding; F:myosin phosphatase activity; F:metal ion binding; C:plasma membrane | receptor protein-tyrosine kinase; protein-serine/threonine phosphatase |
| g26786 | leucine-rich repeat receptor-like serine/threonine/tyrosine-protein kinase SOBIR1 | 279 | F:GO:0004672; C:GO:0016020 | F:protein kinase activity; C:membrane | Transferring phosphorus-containing groups |
| g26789 | leucine-rich repeat receptor-like serine/threonine/tyrosine-protein kinase SOBIR1 | 1011 | P:GO:0018108; F:GO:0004674; F:GO:0004714; F:GO:0005524; C:GO:0005886 | P:peptidyl-tyrosine phosphorylation; F:protein serine/threonine kinase activity; F:transmembrane receptor protein tyrosine kinase activity; F:ATP binding; C:plasma membrane | receptor protein-tyrosine kinase |
| g27161 | putative leucine-rich repeat receptor-like protein kinase | 3339 | P:GO:0006468; F:GO:0004672; F:GO:0005524; C:GO:0016020 | P:protein phosphorylation; F:protein kinase activity; F:ATP binding; C:membrane | Transferring phosphorus-containing groups |
| g27298 | putative inactive leucine-rich repeat receptor-like protein kinase | 2244 | P:GO:0006468; F:GO:0004672; F:GO:0005524; C:GO:0016020 | P:protein phosphorylation; F:protein kinase activity; F:ATP binding; C:membrane | Transferring phosphorus-containing groups |
| g27347 | Acidic leucine-rich nuclear phosphoprotein 32 family B protein | 1302 | C:GO:0016020 | C:membrane | |

| | | | | | |
|---|---|---|---|---|---|
| g27780 | probably inactive leucine-rich repeat receptor-like protein kinase IMK2 | 1185 | F:GO:0005515; C:GO:0016020 | F:protein binding; C:membrane | |
| g3299 | Proline-, glutamic acid- and leucine-rich protein | 357 | | | |
| g5820 | leucine-rich repeat receptor-like serine/threonine-protein kinase BAM1 | 294 | P:GO:0006468; P:GO:0009755; F:GO:0004672; F:GO:0005524; C:GO:0005886; C:GO:0016020 | P:protein phosphorylation; P:hormone-mediated signaling pathway; F:protein kinase activity; F:ATP binding; C:plasma membrane; C:membrane | Transferring phosphorus-containing groups |
| g5822 | leucine-rich repeat receptor-like serine/threonine-protein kinase BAM1 | 2691 | P:GO:0006468; P:GO:0009755; F:GO:0004672; F:GO:0005515; F:GO:0005524; C:GO:0005886; C:GO:0016020 | P:protein phosphorylation; P:hormone-mediated signaling pathway; F:protein kinase activity; F:protein binding; F:ATP binding; C:plasma membrane; C:membrane | Transferring phosphorus-containing groups |
| g6044 | pollen-specific leucine-rich repeat extensin-like protein 3 | 3210 | | | |
| g6113 | putative leucine-rich repeat receptor-like protein kinase | 2772 | P:GO:0006468; F:GO:0004674; F:GO:0005524; C:GO:0016020 | P:protein phosphorylation; F:protein serine/threonine kinase activity; F:ATP binding; C:membrane | Transferring phosphorus-containing groups |
| g618 | leucine-rich repeat protein 2-like | 630 | | | |
| g6345 | leucine-rich repeat protein 1-like | 591 | | | |
| g6533 | probable leucine-rich repeat receptor-like protein kinase At1g35710 | 1932 | P:GO:0016310; F:GO:0005515; F:GO:0016301; C:GO:0005886 | P:phosphorylation; F:protein binding; F:kinase activity; C:plasma membrane | Transferring phosphorus-containing groups |
| g696 | putative leucine-rich repeat receptor-like serine/threonine-protein kinase At2g24130 | 1167 | F:GO:0005515 | F:protein binding | |
| g707 | leucine-rich repeat (lrr) family protein | 1521 | | | |

| | | | | | |
|---|---|---|---|---|---|
| g7919 | putative inactive leucine-rich repeat receptor-like protein kinase | 2175 | P:GO:0046777; F:GO:0004672; F:GO:0005515; F:GO:0005524; C:GO:0005886; C:GO:0016020 | P:protein autophosphorylation; F:protein kinase activity; F:protein binding; F:ATP binding; C:plasma membrane; C:membrane | Transferring phosphorus-containing groups |
| g8141 | putative leucine-rich repeat receptor-like protein kinase | 1905 | F:GO:0005515 | F:protein binding | |
| g822 | pollen-specific leucine-rich repeat extensin-like protein 3 | 2985 | | | |
| g824 | pollen-specific leucine-rich repeat extensin-like protein 3 | 3075 | F:GO:0005515 | F:protein binding | |
| g825 | pollen-specific leucine-rich repeat extensin-like protein 3 | 2400 | | | |
| g8631 | probably inactive leucine-rich repeat receptor-like protein kinase IMK2 | 2514 | P:GO:0006468; F:GO:0004672; F:GO:0005515; F:GO:0005524; C:GO:0016020 | P:protein phosphorylation; F:protein kinase activity; F:protein binding; F:ATP binding; C:membrane | Transferring phosphorus-containing groups |

**Supplementary Table S10** Genes related to citrus acidity based on tBLASTn results in *C. australis* genome

| **Citrus acidity gene** | **Orthologous genes in C. sinensis** | **C. australis_Gene ID** | **chromosome** | **Protein Description** | **Enzyme** | **GO name** |
|---|---|---|---|---|---|---|
| *PH1* | Cs1g20080 | G16480 | 1 | Magnesium-transporting ATPase P-type 1 | P-type Mg(2+) transporter; P-type Ca(2+) transporter; nucleoside-triphosphate phosphatase | P:calcium ion transmembrane transport; P:magnesium ion transmembrane transport; F:P-type calcium transporter activity; F:ATP binding; F:P-type magnesium transporter activity; F:ATP hydrolysis activity; C:plasma membrane; C:intracellular membrane-bounded organelle |
| *PH5* | Cs1g16150 | G24607 | 5 | ATPase 11 plasma membrane-type-related | H(+)-exporting diphosphatase; P-type H(+)-exporting | P:proton export across plasma membrane; F:ATP binding; F:P-type proton-exporting transporter activity; F:ATP hydrolysis activity; C:plasma membrane; C:membrane |

| | | | | | |
|---|---|---|---|---|---|
| | | | | transporter; nucleoside-triphosphate phosphatase | |
| | G14178 | 6 | ATPase 11 plasma membrane-type-related | H(+)-exporting diphosphatase; P-type H(+)-exporting transporter; nucleoside-triphosphate phosphatase | P:proton export across plasma membrane; F:ATP binding; F:P-type proton-exporting transporter activity; F:ATP hydrolysis activity; C:plasma membrane; C:membrane |
| | G14172 | 6 | ATPase 11 plasma membrane-type-related | H(+)-exporting diphosphatase; P-type H(+)-exporting transporter; nucleoside-triphosphate phosphatase | P:proton export across plasma membrane; F:ATP binding; F:P-type proton-exporting transporter activity; F:ATP hydrolysis activity; C:plasma membrane; C:membrane |
| | G14177 | 6 | ATPase 11 plasma membrane-type-related | H(+)-exporting diphosphatase; P-type H(+)-exporting transporter; nucleoside-triphosphate phosphatase | P:regulation of intracellular pH; P:proton transmembrane transport; F:ATP binding; F:P-type proton-exporting transporter activity; F:ATP hydrolysis activity; C:plasma membrane; C:membrane |
| | G14173 | 6 | ATPase 11 plasma membrane-type-related | H(+)-exporting diphosphatase; P-type H(+)-exporting transporter; nucleoside-triphosphate phosphatase | P:proton export across plasma membrane; F:ATP binding; F:P-type proton-exporting transporter activity; F:ATP hydrolysis activity; C:plasma membrane; C:membrane |
| | G20586 | 4 | Plasma membrane ATPase | H(+)-exporting diphosphatase; P-type H(+)- | P:regulation of intracellular pH; P:proton export across plasma membrane; F:ATP binding; F:P-type proton-exporting transporter |

| | | | | | |
|---|---|---|---|---|---|
| | | | | exporting transporter; nucleoside-triphosphate phosphatase | activity; F:ATP hydrolysis activity; C:plasma membrane; C:membrane |
| | G18223 | 4 | Plasma membrane ATPase | H(+)-exporting diphosphatase; P-type H(+)-exporting transporter; nucleoside-triphosphate phosphatase | P:regulation of intracellular pH; P:proton export across plasma membrane; F:ATP binding; F:P-type proton-exporting transporter activity; F:ATP hydrolysis activity; C:plasma membrane; C:membrane |
| | G14171 | 6 | ATPase 11 plasma membrane-type-related | H(+)-exporting diphosphatase; P-type H(+)-exporting transporter; nucleoside-triphosphate phosphatase | P:proton export across plasma membrane; F:ATP binding; F:P-type proton-exporting transporter activity; F:ATP hydrolysis activity; C:plasma membrane; C:membrane |
| | G12167 | 6 | ATPase 11 plasma membrane-type-related | H(+)-exporting diphosphatase; P-type H(+)-exporting transporter; nucleoside-triphosphate phosphatase | P:proton export across plasma membrane; F:ATP binding; F:P-type proton-exporting transporter activity; F:ATP hydrolysis activity; C:plasma membrane; C:membrane |
| | G20845 | 4 | WD REPEATS REGION domain-containing protein | H(+)-exporting diphosphatase; P-type H(+)-exporting transporter; nucleoside-triphosphate phosphatase | H(+)-exporting diphosphatase; P-type H(+)-exporting transporter; nucleoside-triphosphate phosphatase |
| *CitAco3* | g19909 | 4 | Aconitate hydratase | Aconitate hydratase | - |

| | | | | | |
|---|---|---|---|---|---|
| | g22425 | 2 | Aconitate hydratase | Aconitate hydratase 1 | - |
| | g17125 | 1 | Aconitate hydratase 1 | Aconitate hydratase | Aconitate hydratase |
| *CitIDH (NADP-isocitrate_dehydrogenase)* | g5386 | 3 | isocitrate dehydrogenase (NADP) | isocitrate dehydrogenase (NADP(+)) | P:tricarboxylic acid cycle; P:isocitrate metabolic process; P:NADP metabolic process; F:magnesium ion binding; F:isocitrate dehydrogenase (NADP+) activity; F:NAD binding; C:mitochondrion |
| | g11983 | 9 | peroxisomal NADP dependent isocitrate dehydrogenase | isocitrate dehydrogenase (NADP(+)) | - |
| | g21560 | 2 | isocitrate dehydrogenase (NADP) | isocitrate dehydrogenase (NADP(+)) | - |
| *GS* | g10022 | 9 | glutamine synthetase | glutamine synthetase | P:glutamine biosynthetic process; F:glutamate-ammonia ligase activity; F:ATP binding; C:cytoplasm |
| | g12494 | 6 | glutamine synthetase nodule isozyme | glutamine synthetase | - |
| | g7198 | 7 | glutamine synthetase cytosolic isozyme 1-1 | glutamine synthetase | - |
| *GAD* | g22832 | 2 | | | - |
| | g26170 | 5 | | | - |
| *Noemi/AN1* | g27716 | 5 | basic helix-loop-helix transcription factor family protein | | F:DNA binding; F:protein dimerization activity; C:nucleus |
| *PH3* | G13796 | 6 | WRKY transcription factor 44 isoform X1 | | P:regulation of DNA-templated transcription; F:DNA-binding transcription factor activity; F:sequence-specific DNA binding; C:nucleus |

| | | | | | |
|---|---|---|---|---|---|
| *PH4* | G20930 | 2 | R2R3-MYB family transcription factor | | P:cell differentiation; F:transcription cis-regulatory region binding; C:nucleus |

**Supplementary Table S11** Genes related to the synthesis of volatile compounds production in *C. australis* genome

| **Protein name** | **C. australis ID** | **Chromosome** | **Protein description** | **Enzyme name** | **GO name** |
|---|---|---|---|---|---|
| 1. Phosphomevalonate kinase_PMK | g5179 | 3 | phosphomevalonate kinase (peroxisomal) | phosphomevalonate kinase | P:isopentenyl diphosphate biosynthetic process, mevalonate pathway; F:phosphomevalonate kinase activity; F:ATP binding; C:peroxisome |
| 2. Mevalonate kinase_MVK | g4736 | 3 | Amidase 1 | Acting on carbon-nitrogen bonds, other than peptide bonds; mevalonate kinase | P:sterol biosynthetic process; P:phosphorylation; P:isopentenyl diphosphate biosynthetic process, mevalonate pathway; F:mevalonate kinase activity; F:ATP binding; F:hydrolase activity, acting on carbon-nitrogen (but not peptide) bonds, in linear amides; C:cytosol |
| | g21628 | 2 | glucuronokinase 1 | mevalonate kinase | P:isopentenyl diphosphate biosynthetic process, mevalonate pathway; F:mevalonate kinase activity; F:ATP binding; C:cytosol |
| | g26523 | 5 | glucuronokinase 1 | mevalonate kinase | P:phosphorylation; P:isopentenyl diphosphate biosynthetic process, mevalonate pathway; F:mevalonate kinase activity; F:ATP binding; C:cytosol |

| | | | | | |
|---|---|---|---|---|---|
| 3. 3-hydroxy-3-methylglutaryl-CoAsynthase-2_HMGS | g25195 | 5 | Hydroxymethylglutaryl-CoA synthase | hydroxymethylglutaryl-CoA synthase | P:acetyl-CoA metabolic process; P:farnesyl diphosphate biosynthetic process, mevalonate pathway; P:sterol biosynthetic process; F:hydroxymethylglutaryl-CoA synthase activity |
| | g11631 | 9 | Hydroxymethylglutaryl-CoA synthase | hydroxymethylglutaryl-CoA synthase | P:acetyl-CoA metabolic process; P:farnesyl diphosphate biosynthetic process, mevalonate pathway; P:sterol biosynthetic process; F:hydroxymethylglutaryl-CoA synthase activity |
| 4. geranylgeranyl pyrophosphate synthase_chloroplastic_GGPPS | g16883 | 1 | geranylgeranyl pyrophosphate synthase, chloroplastic | geranylgeranyl diphosphate synthase; (2E,6E)-farnesyl diphosphate synthase | P:isoprenoid biosynthetic process; F:farnesyltranstransferase activity; F:geranyltranstransferase activity |
| | g12487 | 6 | geranylgeranyl pyrophosphate synthase 7, chloroplastic-like | Transferring alkyl or aryl groups, other than methyl groups | P:isoprenoid biosynthetic process; F:prenyltransferase activity |
| | g12488 | 6 | Heterodimeric geranylgeranyl pyrophosphate synthase large subunit 1 | Transferring alkyl or aryl groups, other than methyl groups | P:isoprenoid biosynthetic process; F:prenyltransferase activity |
| | g12489 | 6 | geranylgeranyl pyrophosphate synthase 7, chloroplastic-like | Transferring alkyl or aryl groups, other than methyl groups | P:isoprenoid biosynthetic process; F:prenyltransferase activity |
| | g5700 | 3 | Heterodimeric geranylgeranyl pyrophosphate | Transferases | P:isoprenoid biosynthetic process; F:transferase activity |

| | | | | | |
|---|---|---|---|---|---|
| | | | synthase large subunit 1 | | |
| | g5699 | 3 | Heterodimeric geranylgeranyl pyrophosphate synthase large subunit 1 | Transferases | P:isoprenoid biosynthetic process; F:transferase activity |
| | g5698 | 3 | Heterodimeric geranylgeranyl pyrophosphate synthase large subunit 1 | Transferases | P:isoprenoid biosynthetic process; F:transferase activity |
| | g5688 | 3 | Heterodimeric geranylgeranyl pyrophosphate synthase large subunit 1 | Transferases | P:isoprenoid biosynthetic process; F:transferase activity |
| | g5687 | 3 | geranylgeranyl pyrophosphate synthase, chloroplastic-like | Transferases | P:isoprenoid biosynthetic process; F:transferase activity |
| | g5686 | 3 | Heterodimeric geranylgeranyl pyrophosphate synthase large subunit 1 | Transferases | P:isoprenoid biosynthetic process; F:transferase activity |
| | g5684 | 3 | Heterodimeric geranylgeranyl pyrophosphate synthase large subunit 1 | Transferases | P:isoprenoid biosynthetic process; F:transferase activity |
| | g5675 | 3 | Heterodimeric geranylgeranyl pyrophosphate synthase large subunit 1 | Transferases | P:isoprenoid biosynthetic process; F:transferase activity |
| | g5674 | 3 | geranylgeranyl pyrophosphate | Transferases | P:isoprenoid biosynthetic process; F:transferase activity |

| | | | | | |
|---|---|---|---|---|---|
| | | | synthase, chloroplastic-like | | |
| | g5673 | 3 | Heterodimeric geranylgeranyl pyrophosphate synthase large subunit 1 | Transferases | P:isoprenoid biosynthetic process; F:transferase activity |
| | g388 | 8 | Heterodimeric geranylgeranyl pyrophosphate synthase small subunit | Transferring alkyl or aryl groups, other than methyl groups; 2-alkenal reductase [NAD(P)(+)] | P:isoprenoid biosynthetic process; F:prenyltransferase activity; F:2-alkenal reductase [NAD(P)+] activity |
| | g19914 | 4 | heterodimeric geranylgeranyl pyrophosphate synthase small subunit, chloroplastic-like | Transferring alkyl or aryl groups, other than methyl groups | P:isoprenoid biosynthetic process; F:prenyltransferase activity |
| | g12492 | 6 | geranylgeranyl pyrophosphate synthase, chloroplastic | | P:isoprenoid biosynthetic process |
| | g12486 | 6 | Heterodimeric geranylgeranyl pyrophosphate synthase large subunit 1 | Transferring alkyl or aryl groups, other than methyl groups | P:isoprenoid biosynthetic process; F:prenyltransferase activity |
| 5. Farnesyl pyrophosphate synthase_FPPS | g20109 | 4 | Farnesyl pyrophosphate synthase 2 | (2E,6E)-farnesyl diphosphate synthase | P:farnesyl diphosphate biosynthetic process; F:dimethylallyltranstransferase activity; F:geranyltranstransferase activity; C:cytoplasm |
| 06 geranyl_pyrophosphate_synthase_GPPS_Arabidopsis_thaliana | G23478 | 2 | Solanesyl diphosphate synthase 3 | Transferases | P:isoprenoid biosynthetic process; F:transferase activity |
